# APC^Cdh1^-mediated degradation of Cdh1 is necessary for faithful meiotic chromosome segregation in *S. cerevisiae*

**DOI:** 10.1101/2024.07.01.601619

**Authors:** Denis Ostapenko, Mark J. Solomon

## Abstract

The Anaphase-Promoting Complex/Cyclosome (APC/C) is a ubiquitin ligase that promotes the ubiquitination and subsequent degradation of numerous cell cycle regulators during mitosis and in G1. Proteins are recruited to the APC/C by activator proteins such as Cdh1. During the cell cycle, Cdh1 is subject to precise regulation so that substrates are not degraded prematurely. We have explored the regulation of Cdh1 during the developmental transition into meiosis and sporulation in the budding yeast *S. cerevisiae*. Transition to sporulation medium triggers the degradation of Cdh1. Cdh1 degradation is mediated by the APC/C itself in a “*trans*” mechanism in which one molecule of Cdh1 recruits a second molecule of Cdh1 to the APC/C for ubiquitination. Degradation requires an intact glucose-sensing SNF1 protein kinase complex (orthologous to the mammalian AMPK nutritional sensor), which directly phosphorylates Cdh1 on Ser-200 within an unstructured N-terminal region. In the absence of phosphorylation, expression of a Cdh1-S200A mutant is fully stabilized, leading to chromosome instability and loss of viability. We hypothesize that Cdh1 degradation is necessary for the preservation of cell cycle regulators and chromosome cohesion proteins between the reductional and equational meiotic divisions, which occur without the intervening Gap or S phases found in mitotic cell cycles.

## INTRODUCTION

Protein ubiquitination regulates the stability of numerous proteins controlling cell cycle progression and differentiation. During cell growth, protein ubiquitination is carried out by the coordinated action of an E1 (ubiquitin-activating enzyme), multiple E2s (ubiquitin-conjugating enzymes), and two major E3 complexes (ubiquitin ligases): Skp1-Cullin-F-box protein (SCF) and the anaphase-promoting complex or cyclosome (APC/C) (Barford, 2011; Cardozo & Pagano, 2004; Peters, 2006; Pines, 2011; Primorac & Musacchio, 2013; Thornton & Toczyski, 2006). The resulting covalent attachment of poly-ubiquitin chains targets modified proteins for degradation by the 26S proteasome. The evolutionarily conserved APC/C is an essential RING-type ubiquitin ligase composed of a large catalytic core and a substrate-binding activator protein, Cdc20 and Cdh1 in vegetative cells and, in addition, Ama1 in meiotic cells (Barford, 2011; Primorac & Musacchio, 2013; Visintin *et al*, 1997). The APC/C activators both bring protein substrates to the vicinity of the catalytic core and modulate APC/C enzymatic activity. Both Cdc20 and Cdh1 recognize substrates through direct binding to a degradation motif (degron), typically a Destruction box (D-box, RxxLxxxxN), a KEN motif, an ABBA motif, and their derivatives (Di Fiore *et al*, 2015; Glotzer *et al*, 1991; Pfleger *et al*, 2001). Each of these degradation motifs interacts with a corresponding receptor of the activator’s WD-40 domain, and mutations within these degradation motifs or their receptors prevent substrate recruitment and ubiquitination (Davey & Morgan, 2016; He *et al*, 2013; Qin *et al*, 2016). Following substrate binding, Cdc20 and Cdh1 associate with the Apc1, Cdc23 and Cdc27 APC/C subunits through two evolutionarily conserved motifs, the C-box and invariant IR C-terminal tail, with both contacts being essential for the efficient docking of activators and subsequent substrate ubiquitination (Brown *et al*, 2015; Chang *et al*, 2015).

To prevent unscheduled substrate degradation, the activities of Cdh1 and Cdc20 are tightly regulated by numerous post-translational modifications, association with inhibitory factors, and protein stability. During G1 phase, Cdc20 is degraded in an APC/C^Cdh1^-dependent manner and is also inhibited by the spindle assembly checkpoint, which ensures that all chromosomes are properly attached to the mitotic spindle before the onset of mitosis (Foe *et al*, 2011; Foster & Morgan, 2012; Prinz *et al*, 1998; Shirayama *et al*, 1998). Cdh1 is inhibited by phosphorylation by cyclin-dependent protein kinases (CDK), polo-like kinase Cdc5, and the meiosis-specific kinase Ime2 (Crasta *et al*, 2008; Holt *et al*, 2007; Jaquenoud *et al*, 2002; Zachariae *et al*, 1998; Zhou *et al*, 2003). CDK interacts with Cdh1 in a complex, mutually inhibitory manner: Cdh1 inhibits CDK by promoting the ubiquitination and degradation mitotic B-cyclins, whereas CDK-mediated phosphorylation near a nuclear localization signal (NLS) in Cdh1 and near its C-box causes its nuclear export and prevents its interaction with the core APC/C, respectively (Brandeis & Hunt, 1996; Hockner *et al*, 2016; Irniger & Nasmyth, 1997; Jaspersen *et al*, 1999; Schwab *et al*, 1997; Zachariae *et al*., 1998).

In a natural environment, yeast cells are often exposed to various stresses, including nutrient limitation. Under these conditions, diploid yeast cells can undergo a dramatic differentiation from vegetative growth to meiosis and sporulation, culminating in the production of four haploid spores capable of withstanding prolonged periods of starvation or dehydration (Neiman, 2011). The decision to initiate the meiotic program involves the integration of information from numerous signaling pathways, including those sensing limited carbon and nitrogen sources. Ultimately, APC/C-dependent ubiquitination of the Ume6 transcriptional repressor leads to transcriptional activation of *IME1* (*I*nducer of *ME*iosis) and expression of early meiotic genes (Kassir *et al*, 1988; Mallory *et al*, 2007). The ubiquitin ligase activity of APC/C is essential for all stages of meiosis, but its regulation and substrate selectivity are substantially different from that in vegetative cells (Cooper & Strich, 2011). In contrast to mitosis, where mitotic cyclins are completely degraded in G1 phase by APC^Cdh1^, many APC/C substrates persist between meiosis I and meiosis II to prevent DNA re-replication and to maintain chromosome cohesion between the reductional division of meiosis I and the equatorial division of meiosis II. In addition, *S. cerevisiae* expresses a meiosis-specific APC/C activator, Ama1, a Cdh1 homolog that controls ubiquitination of key regulatory proteins (Cooper *et al*, 2000). Later in meiosis, Ama1-mediated ubiquitination promotes the degradation of Cdc20 and other regulatory proteins, including the polo-like kinase Cdc5, mitotic cyclin Clb3, and the major meiosis-specific transcription factor Ndt80 (Okaz *et al*, 2012; Penkner *et al*, 2005; Tan *et al*, 2013; Tan *et al*, 2011).

While the meiotic appearance of APC/C^Ama1^ and the resultant degradation of Cdc20 have been clear for some time, the fate of Cdh1 has been uncertain. We hypothesized that some mitotic substrates of APC/C^Cdh1^ may need to be protected from degradation during meiosis. Here we report that Cdh1 levels decrease upon growth in sporulation-inducing medium. This degradation requires a diploid mating type locus and low glucose levels, which results in the glucose-sensing Snf1 protein kinase phosphorylating Cdh1 on Ser-200. Cdh1 degradation is mediated *in trans* by APC/C^Cdh1^ and is triggered by Ser-200 phosphorylation, though it does not involve any of the known degradation motifs such as a D-box or KEN-box. Failure to degrade Cdh1 during meiosis leads to chromosomal mis-segregation.

## RESULTS

### Transition to sporulation leads to turnover of Cdh1

During vegetative growth of *S. cerevisiae*, Cdh1 undergoes cycles of phosphorylation and de-phosphorylation and binding to regulatory proteins, but its steady-state level remains relatively constant. We tested a range of growth conditions to determine if any affected the stability or steady-state level of Cdh1. The most dramatic effect was observed upon transfer of diploid cells from rich medium (YPD) to sporulation medium (SPM), which induces cells to enter meiosis and to develop into spores as a protective mechanism during poor environmental conditions. The critical components of SPM are a very low concentration of sugar (0.02% raffinose vs the usual 2.0% glucose), 0.3% potassium acetate, and a very low concentration of amino acids. As shown in Figure 1A (*top panel*), Cdh1 levels remained relatively constant when cells were maintained in rich medium but dropped precipitously between six and twelve hours following the shift to sporulation medium. This timing coincides with the degradation of various APC/C^Cdh1^ substrates such as Clb1 and Cdc5 (Visintin *et al*, 2008; Wasch & Cross, 2002) (data not shown).

**Figure 1.**
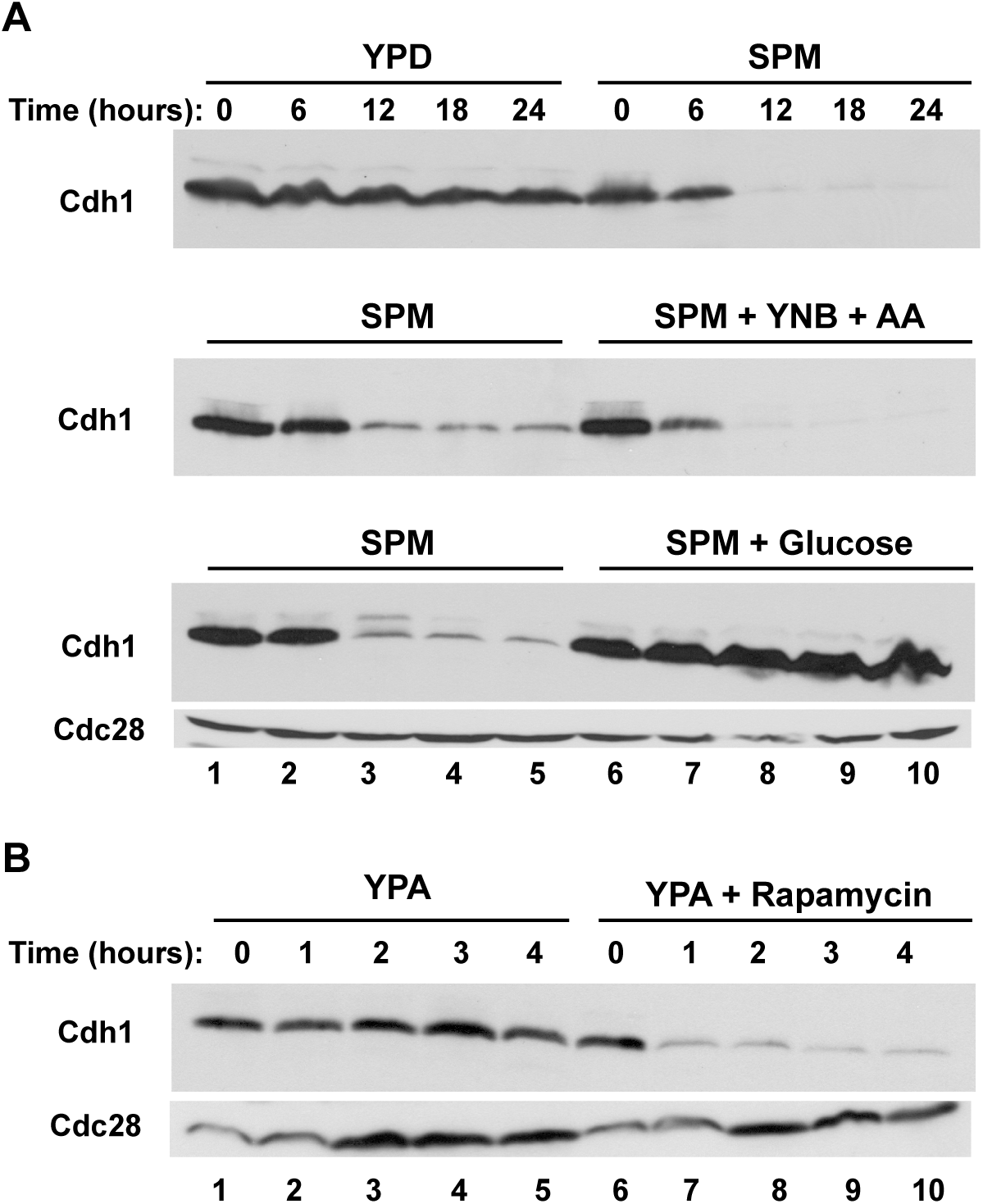
Cdh1 protein levels decline during sporulation. **(A)** *(top)* Cdh1 protein in extracts from an asynchronous population of diploid yeast cells (strain DOY2361) grown in rich medium (YPD, lanes 1-5) or cells transferred from YPA to sporulation medium (SPM, lanes 6-10). Samples were withdrawn at the indicated times and monitored for the presence of Myc-Cdh1 by immunoblotting with 9E10 antibodies. *(middle)* As above with cells transferred to sporulation medium (SPM, lanes 1-5) or to sporulation medium supplemented with 0.67% yeast nitrogen base, 0.2% amino acid mixture (SPM + YNB + AA, lanes 6-10). *(bottom)* As above with cells transferred to sporulation medium (SPM, lanes 1-5) or to sporulation medium supplemented with 1% glucose (lanes 6-10). As a loading control, the membrane was re-probed with anti-PSTAIR antibodies to detect Cdc28. **(B)** Asynchronous cultures of diploid yeast cells (strain DOY2361) pre-grown in acetate-containing medium (YPA) were mock-treated (lanes 1-5) or incubated with 1 µg/ml rapamycin (lanes 6-10). Samples were withdrawn at the indicated times and monitored for the presence of Cdh1 by immunoblotting with anti-Myc antibodies.

Sporulation requires diploidy, a concerted transcriptional program, and low levels of glucose and nitrogen sources. We first explored which nutritional changes were needed for the reduction in Cdh1 level. The addition of amino acids and nitrogen sources to sporulation medium had little effect on the disappearance of Cdh1 (Figure 1A, *middle panel*), suggesting that nitrogen starvation per se was not a required signal. In contrast, addition of 1% glucose (or galactose (not shown)) fully prevented Cdh1 down-regulation (Figure 1A, *bottom panel*). The addition of glucose also prevented the transition to sporulation, which was evident by the absence of tetrads even after prolonged incubation. Thus, a glucose sensing pathway is critical not only for preventing the initiation of the sporulation program but also for the maintenance of Cdh1 protein expression and stability. We will explore the relationship between these events below (Figure 4).

In addition to a nutrient-poor environment, sporulation can be triggered by pharmacological inhibition of the TOR signaling pathway, which mimics starvation for nitrogen and phosphate. Cells grown in YPA (rich in nitrogen sources and acetate, but lacking glucose) maintained their level of Cdh1 (Figure 1B). In contrast, addition of rapamycin caused a rapid (under one hour) degradation of Cdh1. Thus, cells will degrade Cdh1 if they think they’re starving for glucose and nitrogen, even when nitrogen is abundant. We suspect that the more rapid degradation of Cdh1 when cells were transferred from YPA to YPA + rapamycin than when they are transferred from YPD to SPM relates to internal nutrient supplies. Cells switching from YPD to SPM must deplete their internal stores of glucose and excess amino acids. In contrast, cells switching from YPA to YPA + rapamycin have low glucose to begin with and are “tricked” by the presence of rapamycin into thinking they have low nitrogen stores.

In addition to rapamycin, caffeine and methionine sulfoximine (MSX), two chemically unrelated compounds that also inhibit the TOR signaling pathway, also led to decreased Cdh1 levels, whereas other stress conditions, such as high osmolarity, heat shock, oxidative stress, and DNA damage, had no effect (data not shown).

### Diploidy and the initiation of meiosis are not required for Cdh1 down-regulation

Because the experiments in Figure 1 used diploid *MATa/MATα* cells, we wondered whether the degradation of Cdh1 in sporulation medium is a general response to starvation, or if it requires diploidy and the initiation of the meiotic program. Natural population of *S. cerevisiae* exist in three developmentally distinct mating types, which are distinguished by the expression of transcription factors from their mating-type loci. Haploids contain *MATa* or *MATα* at their mating-type locus whereas diploids are heterozygous for this locus (*MATa/MATα*). Diploid cells can undergo meiosis and sporulation to produce haploid spores, which can germinate to produce haploid cells capable of mating with cells of the opposite mating type to re-form diploids.

Although Cdh1 was unstable in *MATa/MATα* grown in sporulation medium (Figure 2A, *top panel, left*), Cdh1 was fully stable in *MATa* haploids (Figure 2A, *top panel, right*), which arrested as individual unbudded cells rather than proceeding though sporulation. Since *MATa/MATα* cells differ from *MATa* cells in both ploidy and mating type, we tested Cdh1 stability in *MATa/MATa* diploids to distinguish which factor was important. *MATa/MATa* diploids have the ploidy of *MATa/MATα* cells but have the mating type of *MATa* cells. Interestingly, Cdh1 was fully stable in *MATa/MATa* diploid cells shifted to sporulation medium (Figure 2A, *middle panel*), indicating that it is mating type, and not ploidy, that is important for the decline in Cdh1 levels. Finally, we found that addition of the *α2* transcription factor to a *MATa* haploid, thereby converting these cells to the *MATa/MATα* transcriptional profile, led to Cdh1 turnover (Figure 2A, *lower panel*). Therefore, Cdh1 down-regulation requires the transcription profile of *MATa/MATα* cells but does not require diploidy *per* se.

**Figure 2.**
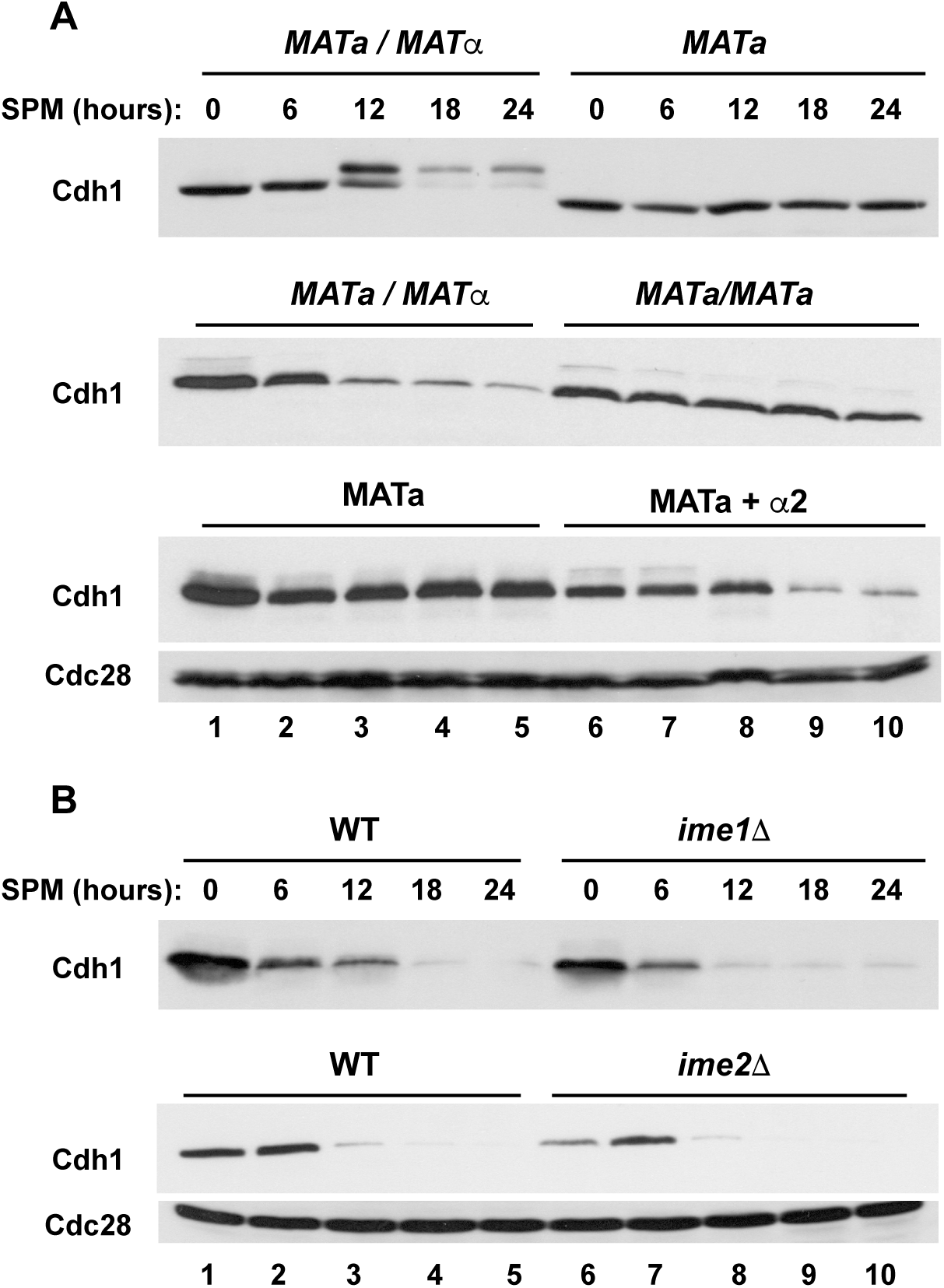
Cdh1 degradation depends on heterozygosity of the mating type locus but not on cell ploidy per se. **(A)** *(top)* Asynchronous diploid *MATa/MATα* cells (strain DOY2361) (lanes 1-5) and haploid *MATa* cells (strain DOY2282) (lanes 6-10) were transferred from YPA to sporulation medium as in Figure 1A. Samples were withdrawn at the indicated times and processed for immunoblotting to detect Myc-tagged Cdh1. *(middle)* The same time courses as above were performed using two isogenic diploid strains that differ only in the configuration at the mating-type loci: *MATa/MATα* (strain DOY2780) (lanes 1-5) and *MATa/MATa* (strain DOY2783) (lanes 6-10). Only the *MATa/MATα* cells were able to sporulate and produce tetrads. *(bottom)* The same time courses were performed using a haploid *MATa* strain carrying either an empty vector (lanes 1-5) or a vector expressing the α2 transcription factor (strain DOY2786) normally expressed in *MATa/MATα* diploid cells, thus making these cells equivalent to *MATa/MATα* cells at the mating-type locus but otherwise haploid (lanes 6-10). The *MATa + α2* cells could not undergo sporulation. Anti-Cdc28 probing of the same filter was used as a loading control. **(B)** Expression of Inducer of MEiosis genes is not required for Cdh1 degradation. Homozygous diploid strains that were wild-type (strain DOY2361) or deleted for *IME1* (strain DOY2554), or *IME2* (strain DOY2555) were transferred to sporulation medium, and tested at the indicated times for Cdh1 expression. Mutation of either *IME* gene prevents transcription of early meiotic genes and yeast sporulation. Samples were processed for immunoblotting with 9E10 antibodies to detect Myc-Cdh1 protein. The membrane was re-probed with anti-PSTAIR antibodies to detect Cdc28 as a loading control.

Although we have shown that conditions that lead to meiosis and sporulation (diploidy and starvation) also lead to reduced levels of Cdh1, we have not shown that entry into meiosis is necessary for the decline in Cdh1 level. To do so, we examined Cdh1 levels in two independent *IME* (*I*nitiator of *ME*iosis) mutants. *IME1* encodes the major transcriptional activator of early meiotic genes. Its promoter harbors numerous cis-acting sensors for cell ploidy, glucose, and nitrogen levels that together determine whether external conditions are appropriate to begin the transition into meiosis and sporulation. *IME2* encodes a Cdc28-like protein kinase that phosphorylates multiple targets, including Cdh1, and is a direct transcriptional target of Ime1. In the absence of *IME* genes, cells fail to enter the meiotic program and continue in the vegetative state. Remarkably, deletion of either of these meiotic inducers had no effect on the decline of Cdh1 (Figure 2B, lanes 6-10). Thus, the down-regulation of Cdh1 is signaled prior to or in parallel to the induction of meiotic entry by the *IME* genes, but does not require the initiation of the meiotic transcriptional program.

### APC/C^Cdh1^-dependent degradation of Cdh1 *in trans*

Before determining which signaling pathway might be responsible for the decline in Cdh1 level, we first verified that the decrease in Cdh1 is ubiquitin-dependent. Addition of the proteasome inhibitor MG-132 blocked the decline in Cdh1 upon transfer to sporulation medium (Figure 3A). The gradual increase in the amount of Cdh1 is presumably due to new synthesis in the absence of degradation.

**Figure 3.**
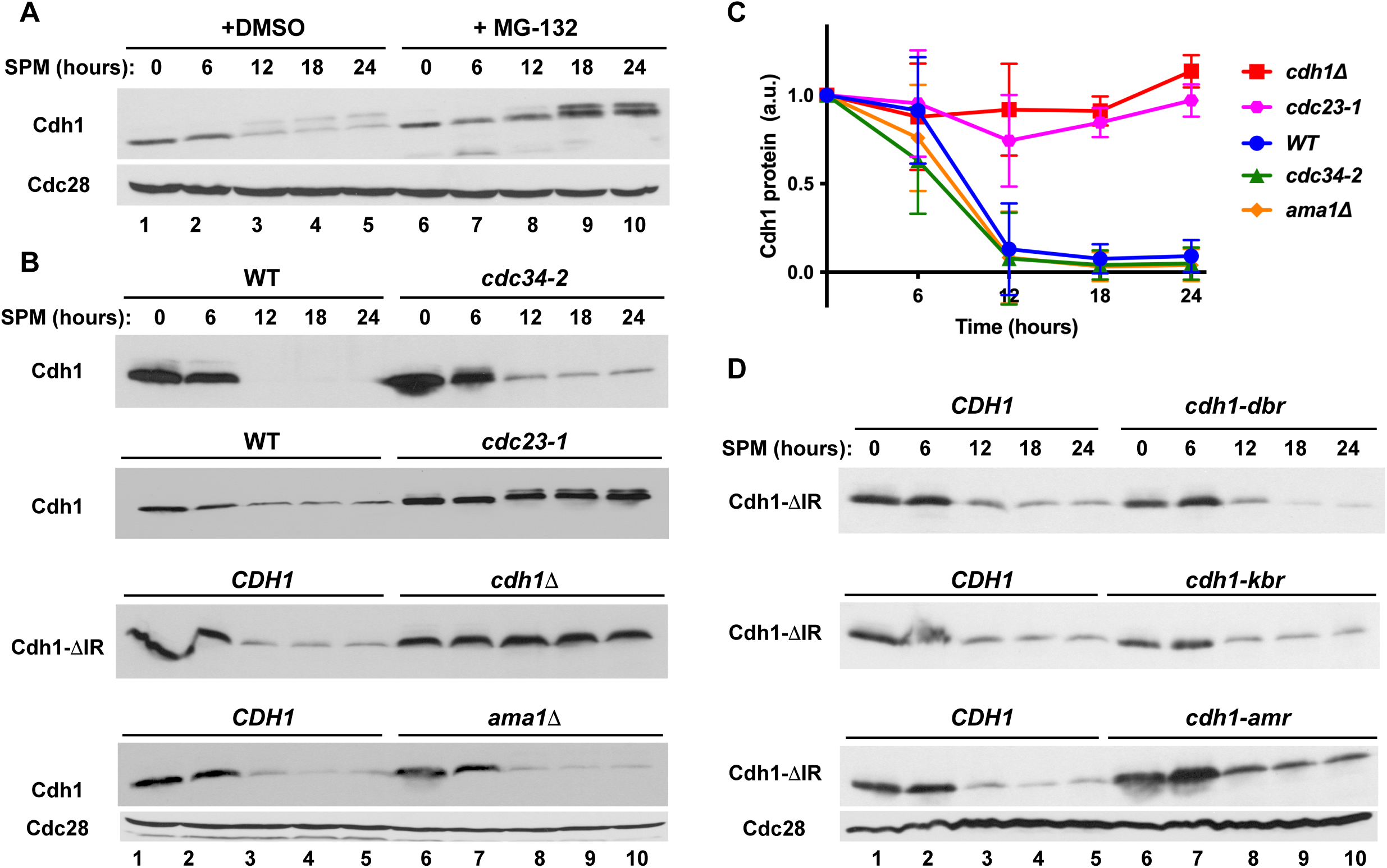
Cdh1 is degraded in an APC/C-dependent manner during sporulation. **(A)** *pdr5Δ* homozygous diploid cells (which are permeable to the proteasome inhibitor MG132; strain DOY2989) were transferred to sporulation medium in the absence (lanes 1-5) or presence of 20 μM MG132 to inhibit the proteasome. Samples were withdrawn at the indicated times and processed to detect Myc-Cdh1 as in Figure 1A. Anti-Cdc28 was used as a loading control. **(B)** Wild-type and isogenic homozygous diploid strains carrying a conditional allele of the ubiquitin-conjugating enzyme of the SCF complex (*cdc34-2, top panel*; strain DOY2542), or an APC/C core subunit (*cdc23-1, second panel*; strain DOY2540) were grown at permissive conditions and transferred to sporulation medium at 35°C to inactivate the temperature-sensitive proteins. Of note, transfer to 35°C did not interfere with the ability of wild-type cells to form tetrads but arrested cell cycle progression of the *cdc34-2* and *cdc23-1* strains. (*Third panel*) Wild type *CDH1* (strain DOY2719) and *cdh1Δ* mutant (strain DOY2725) strains carried an endogenous *Myc-CDH1-ΔIR* (“Cdh1-ΔIR”) allele, which lacks the C-terminal two amino acid residues of Cdh1 necessary for binding to the APC/C core. Therefore, the Cdh1-ΔIR protein is non-functional in promoting the degradation of APC/C substrates, but can serve as a substrate itself. Samples were processed to detect Myc-Cdh1-ΔIR protein levels as in Figure 1A. (*Bottom panel*) A wild-type and an isogenic homozygous diploid *ama1Δ* strain (strain DOY2602), which carries a homozygous deletion of the meiosis-specific APC/C activator, were transferred to sporulation-inducing medium and incubated at 30°C for the indicated times. Samples were processed to detect Myc-Cdh1-ΔIR protein levels as in Figure 1A. **(C)** The relative stability of Cdh1 in wild-type and the indicated mutant strains was quantified by ImageJ software (http://www.ImageJ.net). The initial level of Cdh1 in each individual strain at time 0 was set to 100%. Data represent the means and standard deviations from three independent experiments. **(D)** All strains expressed the non-functional “Cdh1-ΔIR” reporter described in (C) and either a wild-type copy of *CDH1* (strain DOY2970;), or a form of Cdh1 that has been mutated so that it can no longer recognize Destruction Boxes (*cdh1-dbr*; strain DOY2971; *top*), KEN boxes (*cdh1-mkr*; strain DOY2972; *middle*), or the ABBA motif (*cdh1-amr*; strain DOY2973; *bottom*). All strains were transferred to sporulation medium and incubated for the indicated times. Samples were withdrawn and immunoblotted with 9E10 antibodies to detect Myc-Cdh1-ΔIR. Immunoblotting with an anti-Cdc28 antibody was used as a loading control.

In considering which ubiquitin ligase might be responsible for Cdh1 degradation, we first turned to SCF (Skp1-Cullin-F box protein) complexes since they typically target phosphorylated proteins; Cdh1 is known to be phosphorylated at multiple sites and the transition to sporulation coincides with the appearance of a reduced-mobility form of Cdh1 upon SDS-PAGE (for example, see Figure 3A). We shifted wild-type or *cdc34-*2 temperature-sensitive mutant cells into sporulation medium at the non-permissive temperature (35°C) and followed Cdh1 levels. Cdc34 is a ubiquitin-conjugating enzyme (E2) that functions with all SCF complexes in yeast. We noticed that wild-type cells were able to sporulate and formed tetrads at the elevated temperature (35°C), whereas *cdc34-2* mutant arrested as single cells, likely due to abnormally high levels of multiple SCF substrates. Even though *cdc34-2* cells did not proceed through meiosis, Cdh1 was still down-regulated to the same extent as in wild-type cells (Figure 3B, *top panel,* and Figure 3C). Similarly, inactivation of Cdc4, a major F-box protein for the SCF, also failed to stabilize Cdh1 (data not shown).

We next asked whether the APC/C itself could promote Cdh1 degradation. We envisioned that Cdh1 might transition from being an APC/C activator to being an APC/C substrate. Two examples of such a transition are well documented in vegetative cells, first, in which APC/C^Cdc20^ promotes the ubiquitination of the Cdc20 in the APC/C complex (“*in cis*”) during mitosis, and second, in which APC/C^Cdh1^ promotes the ubiquitination of Cdc20 (“*in trans*”) in G1 (Foe *et al*., 2011; Robbins & Cross, 2010). We tested whether the APC/C was involved in Cdh1 degradation by transferring wild-type cells and *cdc23-1* cells (containing a temperature-sensitive mutation in a core APC/C subunit) into sporulation medium at the non-permissive temperature (35°C) and following Cdh1 levels. The elevated temperature did not affect Cdh1 degradation in wild-type cells but it completely blocked degradation in the *cdc23-1* cells (Figure 3B, *second panel*), indicating that the APC/C is involved in Cdh1 degradation.

Having determined that the APC/C core complex was essential for Cdh1 turnover, we next asked which activator protein was involved in this process. Ama1 is a meiosis-specific homolog of Cdc20 and Cdh1 that promotes the ubiquitination of multiple regulatory proteins, including Cdc20. Since *AMA1* is not essential for vegetative growth, we examined Cdh1 levels in *ama1Δ* mutant cells after their transition to sporulation medium. As expected, *ama1Δ* cells were unable to enter meiosis. However, Cdh1 turnover occurred as well in *ama1Δ* cells as in wild-type cells (Figure 3B, *bottom panel*), indicating that Ama1 was not required for Cdh1 turnover.

Excluding Ama1 raised the possibility that Cdh1 might promote its own degradation. One version of such auto-ubiquitination (termed “*trans*”) involves two molecules of Cdh1, one occupying the conventional activator position in an APC/C^Cdh1^ complex, and the second serving as the substrate that is bound to the APC/C^Cdh1^ complex. To test this possibility, we expressed a form of Cdh1 termed Cdh1-ΔIR, which lacks its C-terminal two amino acids and is cannot interact with the APC/C core subunit Cdc23. It is thus unable to serve as an APC/C activator, but could potentially serve as a substrate. (Preliminary testing in haploid cells showed that Cdh1-ΔIR was unable to support the degradation of the Clb2 APC/C substrate (data not shown).) When cells expressing Cdh1-ΔIR but no other form of Cdh1 were transferred to sporulation medium, Cdh1-ΔIR was stable (Figure 3A, *third panel*, lanes 6-10, and Figure 3C). However, when the cells also expressed wild-type Cdh1, Cdh1-ΔIR was degraded with normal kinetics (Figure 3A, *third panel*, lanes 1-5, and Figure 3C), indicating that it was degraded *in trans*.

Although most APC/C substrates and other Cdc20-or Cdh1-binding proteins contain one of the common degradation motifs (a Destruction Box, a KEN box, or, less commonly, an ABBA motif), Cdh1 itself does not contain an obvious version of any of these motifs. Some APC/C substrates contain a degenerate form of one of these motifs that binds to the same binding surface on Cdc20 or Cdh1 as conventional substrates. Structural studies have revealed the ‘receptors’ within the WD-40 domains of the APC/C activators that are specific for the three different types of APC/C degrons. We determined whether Cdh1 might contain a “cryptic” degradation motif by testing whether Cdh1-ΔIR was stabilized in the presence of otherwise wild-type Cdh1 containing one of the receptor mutants. Remarkably, Cdh1-ΔIR was degraded with wild-type kinetics in the presence of Cdh1 with a Destruction Box receptor (*cdh1-dbr*), a KEN box receptor (*cdh1-kbr*), or an ABBA receptor (*cdh1-amr*) (Figure 3D) even though these mutants were deficient in targeting most of the known substrates in G1-arrested cells (Qin *et al*., 2016). Thus, Cdh1-Cdh1 recognition during the sporulation program involves a non-canonical degradation motif on the substrate Cdh1 and a novel binding surface on the activator Cdh1. A deletion analysis of Cdh1 suggests that a novel degron may be located between amino acids 50 and 150 (Supplemental Figure 3).

### Cdh1 down-regulation is mediated by the SNF1 protein kinase complex

Multiple pathways sense glucose levels in budding yeast. Glucose starvation inhibits protein kinase A (PKA) and thereby induces the sporulation pathway. However, deletion or inhibition of all three PKA catalytic subunits had no effect on Cdh1 degradation in sporulation medium (Supplemental Figure 4A, *top panel*). Likewise, deletion of two other proteins involved in nutrient signaling and the initiation of meiosis and sporulation (Rim11 and Mck1) also had no impact on Cdh1 down-regulation (Supplementary Figure 4A, *bottom panel*).

Another major pathway for sensing the presence of glucose involves the SNF1 protein kinase complex. The SNF1complex (ortholog of the mammalian AMPK nutrient sensor) is required for yeast cells to survive in the absence of fermentable glucose and other environmental stresses. SNF1 protein kinase activity increases in the presence of alternative carbon sources and during sporulation but is rapidly inhibited in the presence of glucose. Thus, in the absence of Snf1, SNF1-regulated pathways are unaffected by the absence of glucose. Notably, deletion of *SNF1* prevented down-regulation of Cdh1 in sporulation medium (Figure 4A, *top panel*, and Figure 4B). Consistent with their functioning in a common pathway, both SNF1 and CDH1 are essential for survival under nutrient-limiting conditions (Figure 4D). The SNF1 complex also contains one of three potential β subunits, which differ in their subcellular localization and substrate specificity, and a unique γ-subunit, Snf4. None of the single β subunit deletions (*GAL83, SIP1,* and *SIP2*) affected Cdh1 levels (Supplemental Figure 4, top three *panels*), indicating functional redundancy among these proteins. In contrast, deletion of the sole γ-subunit, *SNF4*, fully stabilized Cdh1 (Figure 4A, *second panel*, and Figure 4B).

**Figure 4.**
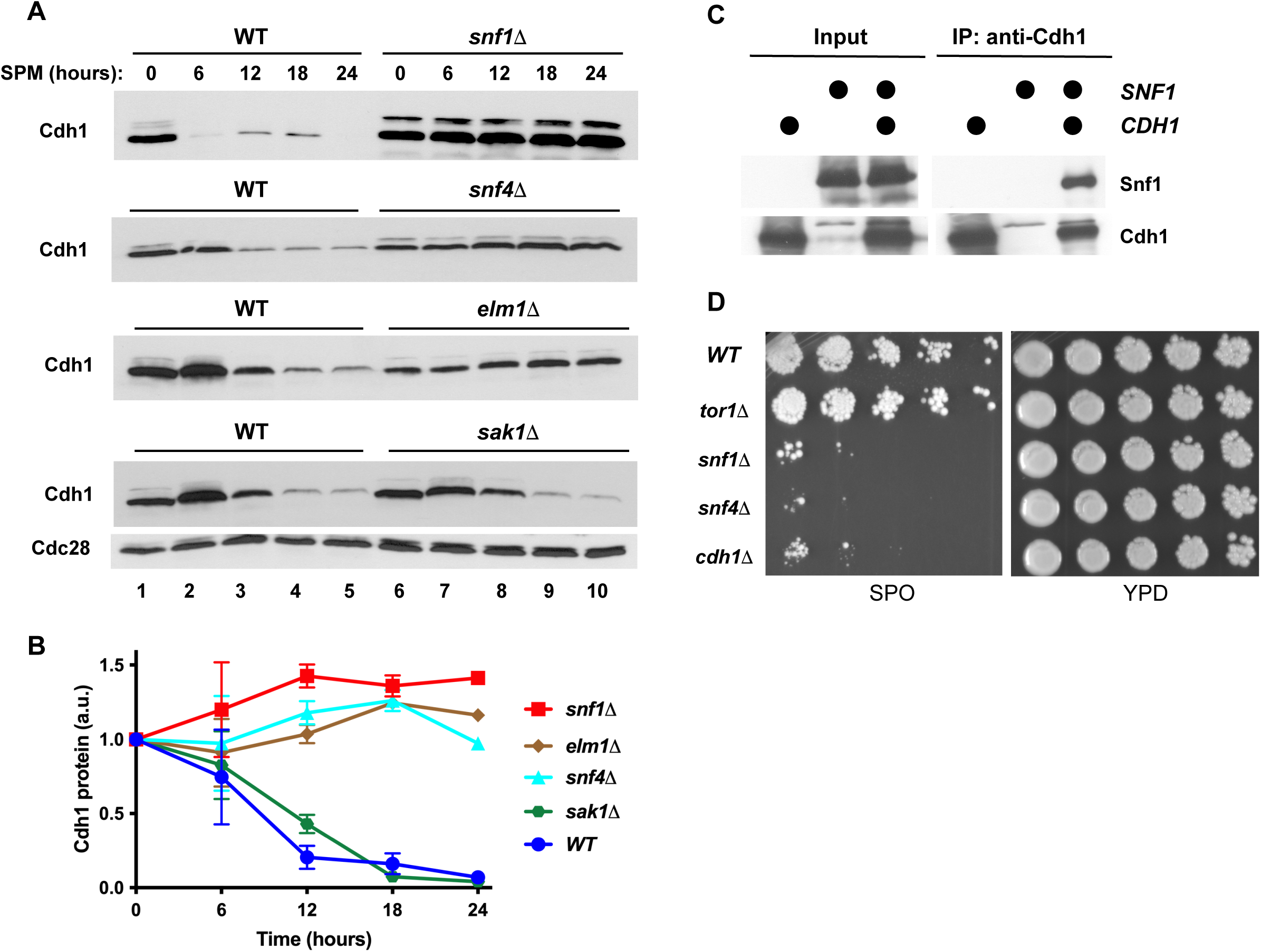
The glucose-sensing AMP-dependent protein kinase (AMPK) pathway is required for Cdh1 turnover and cell survival during sporulation. **(A)** A wild-type strain (strain DOY2361) and isogenic mutant strains carrying homozygous deletions of the AMPK/SNF1 alpha subunit (*snf1Δ* strain DOY2603), the SNF1 regulatory subunit (*snf4Δ* strain DOY2655), or the Snf1-activating kinases Elm1 (*elm1Δ* strain DOY2676) or Sak1 (*sak1Δ* strain DOY2675) were grown in YPA and transferred to sporulation medium. Samples were processed to detect Myc-Cdh1 protein as in Figure 1A. **(B)** The relative abundance of Cdh1 in the wild-type strain and in the indicated mutant strains in (A) were quantified using ImageJ software and normalized relative to the level of Cdc28 protein from the same sample. The initial amount of Cdh1 at time 0 was set at 100% and the fractions of the protein remaining in the samples at subsequent times were plotted relative to this starting level. **(C)** Cdh1 interacts with SNF1 *in vivo*. Yeast strains carrying endogenous *SNF1-TAP* and galactose-inducible *GST-CDH1* and *GST* expression vectors, as indicated on the figure, were activated by galactose for four hours. GST-Cdh1 was immunoprecipitated from cell extracts with glutathione beads and probed for the presence of Snf1-TAP by immunoblotting with PAP antibodies. The levels of GST-Cdh1 was determined by immunoblotting anti-Cdh1 antibodies. **(D)** Wild type (strain DOY2361) and three isogenic diploid strains carrying homozygous deletions of *TOR1* (strain DOY2512), *SNF1* (strain DOY2603), *SNF4* (strain DOY2655), and *CDH1* (strain DOY2589) were pre-incubated in sporulation-inducing medium (SPO) for 7 days at 30°C, serially diluted, and spotted on a plate containing rich medium (YPD). Asynchronous control cultures of the same strains grown in rich medium (YPD) were diluted and plated in the same order; both plates were grown at 30°C for three days.

Activation of the Snf1 protein kinase requires binding to Snf4 as well as phosphorylation at Thr270 by one of three upstream protein kinases: Sak1, Elm1, and Tos3. The activities of these partially redundant protein kinases are ultimately linked to intracellular AMP concentration, which in turn depends on the availability of glucose. We tested whether deletion of any of these upstream protein kinases affected Cdh1 levels. Surprisingly, deletion of *ELM1* fully stabilized Cdh1, resembling the effects observed with *snf1Δ* and *snf4Δ* mutant strains (Figure 4A, *third panel*, and Figure 4B). However, deletion of *SAK1* had negligible effects on Cdh1 stability (Figure 4A, *bottom panel*, and Figure 4B), even though its abundance and overall activity are significantly higher than the other two upstream kinases (Ben Turk, Yale University, personal communication). Deletion of *TOS3* also had no effect on Cdh1 levels (data not shown). Thus, of the three protein kinases directly upstream of Snf1, Elm1 acts uniquely to activate Snf1 in a pathway from glucose limitation to Cdh1 down-regulation. However, it should be noted, that in addition to Snf1, Elm1 phosphorylates numerous other protein targets, so we cannot exclude the possibility that the stabilizing effect of *elm1Δ* might occur through multiple pathways.

SNF1 exerts its major effects through large-scale reprogramming of gene transcription in response to glucose depletion. For example, Snf1 directly phosphorylates the Mig1 transcriptional repressor, which leads to its release from gene promoters required for utilization of alternative carbon sources. Snf1 also phosphorylates the transcriptional activator Adr1, leading to the transcription of many metabolic genes, and the N-terminal domain of histone H3, which leads to transcriptional changes within hundreds of genes. To examine the possibility that increased Cdh1 turnover is caused by changes in gene transcription downstream to *SNF1*, we examined Cdh1 protein levels in *mig1Δ* and *adr1Δ* mutant strains. We found that neither deletion affected Cdh1 levels--in sharp contrast to the effect in *snf1Δ* and *snf4*Δ cells--making it unlikely that expression of one of the many Mig1-repressed or Adr1-activated genes is responsible for Cdh1 instability (Supplemental Figure 4B, *bottom two panels*). These negative results raised the possibility that SNF1 might regulate Cdh1 stability directly; this hypothesis was supported by the finding that GST-Cdh1 could pull down Snf1 from cell extracts (Figure 4C).

### Snf1 phosphorylates Cdh1 on Ser-200

Because Cdh1 is regulated via phosphorylation during vegetative growth, we wondered whether phosphorylation might affect its stability during meiosis. To address this question, we purified Cdh1 from diploid yeast cells grown in nutrient-rich medium (YPG) and from the same cells grown in sporulation medium (SPO) for six hours, at which point Cdh1 resolves as two bands on SDS-PAGE. We compared the composition and relative abundance of individual phospho-peptides in the two samples by phospho-proteomic mass spec analysis (Figures 5A and 5B). Along with decreased phosphorylation of multiple CDK phosphorylation sites (open bars), we noted phosphorylation of three non-CDK sites (Ser32, Ser50 and Ser-200, solid bars) that partially fit the SNF1 consensus recognition sequence, ΦxR/KxxS/TxxxΦ (where Φ represents a hydrophobic amino acid) (Treitel *et al*, 1998).

Based on these findings, we tested whether Snf1 could phosphorylate Cdh1 on one or more of these sites *in vitro*. We purified constitutively-active Snf1-G53R (Estruch *et al*, 1992) from diploid cells, and incubated it with a purified N-terminal fragment of Cdh1 (Cdh1-NTD). The reaction products were separated on a Phos-tag^TM^ SDS-PAGE gel, which enhances the mobility shifts resulting from protein phosphorylation (Kinoshita *et al*, 2006). Sfn1 phosphorylated Cdh1-NTD in an ATP-dependent manner, as evident from the reduced mobility of Cdh1-NTD (Figure 5D, *top panel, lanes 1-4*). Mutation of Ser-200 to Ala-200 abolished the mobility shift (Figure 5D, lanes 9 and 10), whereas mutations of Ser-32 and Ser-50 to alanines had no effect on Cdh1-NTD phosphorylation (Figure 5D, lanes 5-8), indicating that Ser-200 is the major Snf1 phospho-acceptor site within Cdh1-NTD.

**Figure 5.**
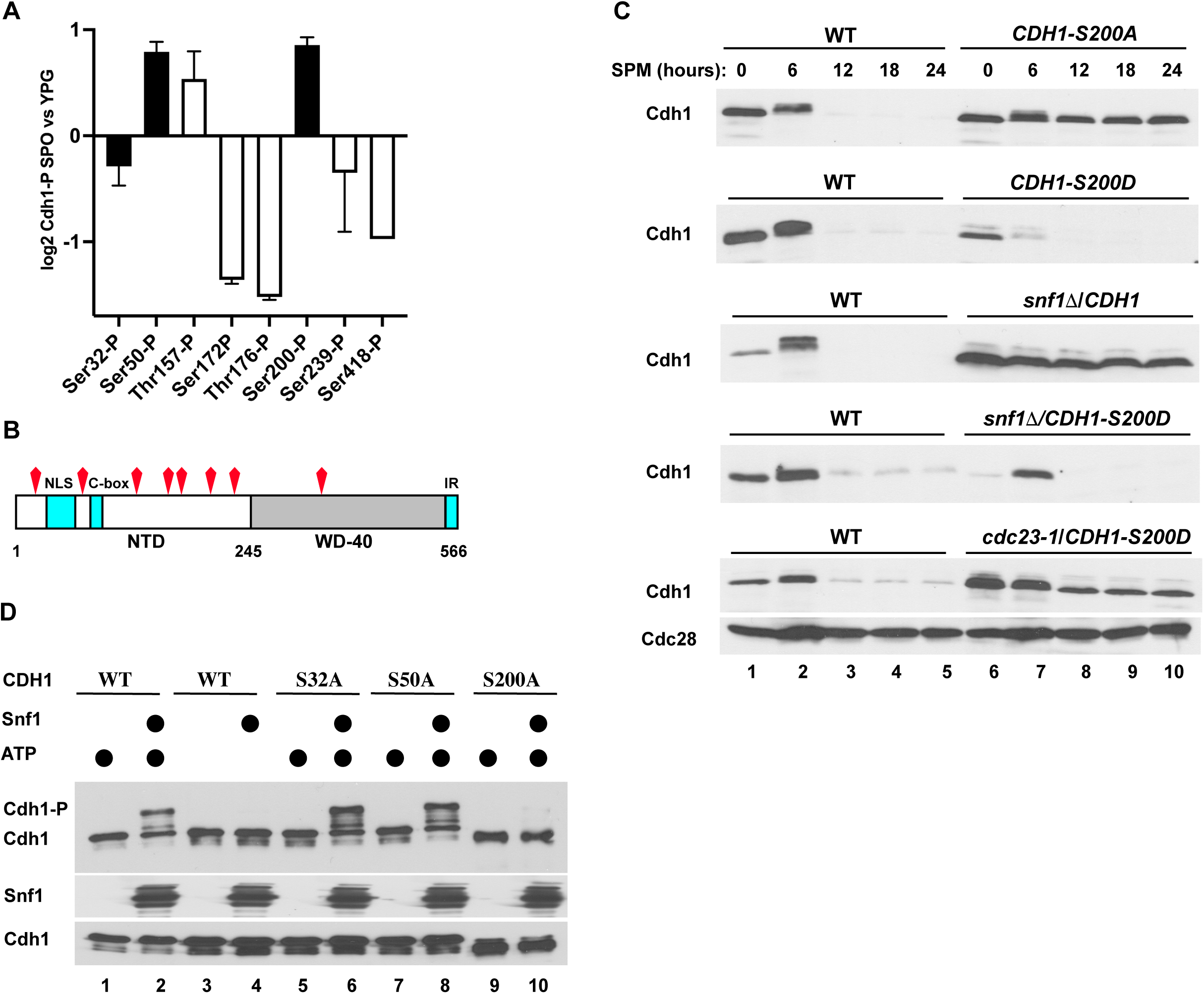
Snf1-mediated phosphorylation regulates Cdh1 stability. **(A)** Wild-type diploid cells carrying *GST-CDH1* (strain DOY3384) were grown in YP-galactose (YPG) at 30°C or transferred to sporulation medium (SPO) for 6 hours at 30°C. GST-Cdh1 was affinity purified and subjected to mass-spectrometry to detect phospho-peptides. The phosphorylated peptides corresponding to indicated Cdh1 sites were quantified using Scaffold5 software (https://www.proteomesoftware.com), normalized relative to the total peptide counts, log2 transformed, and compared between SPO and YPG cultures. Open bars within the graph represent known CDK phosphorylation sites, and closed bars indicate sites that resemble the SNF1 consensus phosphorylation site. **(B)** A cartoon of Cdh1 indicating the phosphorylation sites shown in (A). Most of the differentially-phosphorylated sites are located within the unstructured N-terminal domain (NTD), including a cluster of six CDK sites (Thr157, Ser169, Thr173, Thr176, Ser227, Ser239). Of note, peptides corresponding to three other CDK sites (Thr12, Ser16, and Ser42) were not detected by mass-spectrometry. The second cluster of phospho-sites (Ser32, Ser50, Se200) is located within the region that regulates Cdh1 localization and its interaction with the core APC (Hockner *et al*., 2016). The locations of an NLS (nuclear localization signal), the C-box, and the IR motifs are indicated. **(C)** Phosphorylation of Ser-200 regulates Cdh1 stability during meiosis. Ser-200 was mutated to either alanine or aspartic acid. *(top two panels)* Wild type *CDH1* (strain DOY2899) and two heterozygous diploid strains, *CDH1-S200A* (strain DOY4043) and *CDH1-S200D* (strain DOY4011) (carrying an untagged copy of wild-type *CDH1*) were tested after shift to sporulation medium as in Figure 1A. *(third and fourth panels) snf1Δ CDH1* (strain DOY3459) and *snf1Δ CDH1-S200D* (strain DOY4244) strains were grown and analyzed for Myc-Cdh1 expression. *(bottom) cdc23-1 CDH1-S200D* cells (strain DO4246) were pre-grown under permissive conditions and transferred to sporulation medium at 35°C to inactivate Cdc23. Samples were collected at the indicated times and processed to detect Myc-Cdh1 protein as in Figure 1A. **(D)** Snf1 phosphorylates Cdh1 Ser-200 *in vitro*. Constitutively active *GST-SNF1-G53R* was expressed in yeast cells grown in YP-galactose (YPG) (strain DOY4195). The SNF1 protein kinase complex was affinity purified and incubated with recombinant His_6_-Cdh1^1-250^ in the absence and presence of ATP, as indicated. Three putative Snf1 consensus sites (Ser32, Ser50, Ser-200) were individually mutated to alanine. The products of the assay were resolved on a Phos^TM^-tag gel *(top panel),* conventional SDS-PAGE *(bottom panel)* and probed with anti-Cdh1 antibodies. Snf1 was detected by immunoblotting with anti-GST antibodies *(middle panel)*.

### Ser-200 phosphorylation is necessary for Cdh1 degradation during meiosis

Since deletion of *SNF1* stabilizes Cdh1 in sporulation medium, and since Snf1 phosphorylates Cdh1 on Ser-200, we asked whether phosphorylation of Ser-200 is necessary for Cdh1 degradation during meiosis. Whereas wild-type Cdh1 was degraded between 6 and 12 hours of incubation in sporulation medium, Cdh1-S200A was fully stable through the 24-hour time course (Figure 5C, *top panel*), mimicking the stabilization of wild-type Cdh1 in the presence of glucose and in *snf1Δ* cells (Figures 1A and 4A). It is worth noting that *CDH1-S200A* cells progressed through meiosis and formed tetrads, though with a reduced efficiency (see below), in sharp contrast to *snf1Δ* cells that were unable to sporulate. Conversely, Cdh1-S200D, which mimics phosphorylation of this site, was highly unstable (Figure 5C, *second panel*), even in the absence of *SNF1* (Figure 5C, *fourth panel*), indicating that phosphorylation of Ser-200 is the only function of Snf1 in promoting Cdh1 degradation. Degradation of Cdh1-S200D was still dependent on the APC/C (Figure 5C, *bottom panel*). Importantly, the stabilization of Cdh1-S200A is not due to gross misfolding since the mutant could still promote the degradation of APC/C substrates Clb2 and Cdc5 in G1-arrested haploid cells (Supplemental Figure 5B, *lanes 5 and 6*), as well as in diploid cells during sporulation (data not shown). In contrast, Cdh1-S200D was inactive in G1 (Supplemental Figure 5, *lanes 7 and* 8) yet was still rapidly degraded via the APC/C.

### Stabilization of Cdh1 leads to chromosome instability and reduced spore viability

The complete and tightly regulated degradation of Cdh1 during meiosis and sporulation suggests that its elimination might be important for the faithful completion of these processes. We addressed this idea by expressing wild-type, non-degradable (S200A) and constitutively degradable (S200D) versions of Cdh1 in cells placed under sporulation conditions. We observed that wild-type and *S200A* cells incubated in sporulation medium for 14 days grew fine after plating on rich medium (Figure 6A). In contrast, *S200D* cells failed to grow when plated back onto rich medium (Figure 6A), indicating that the proper timing of Cdh1 degradation is important for passage through meiosis and sporulation. Indeed, *cdh1Δ* cells, like *S200D* cells, fail to undergo meiosis and sporulation (Figure 4D).

**Figure 6.**
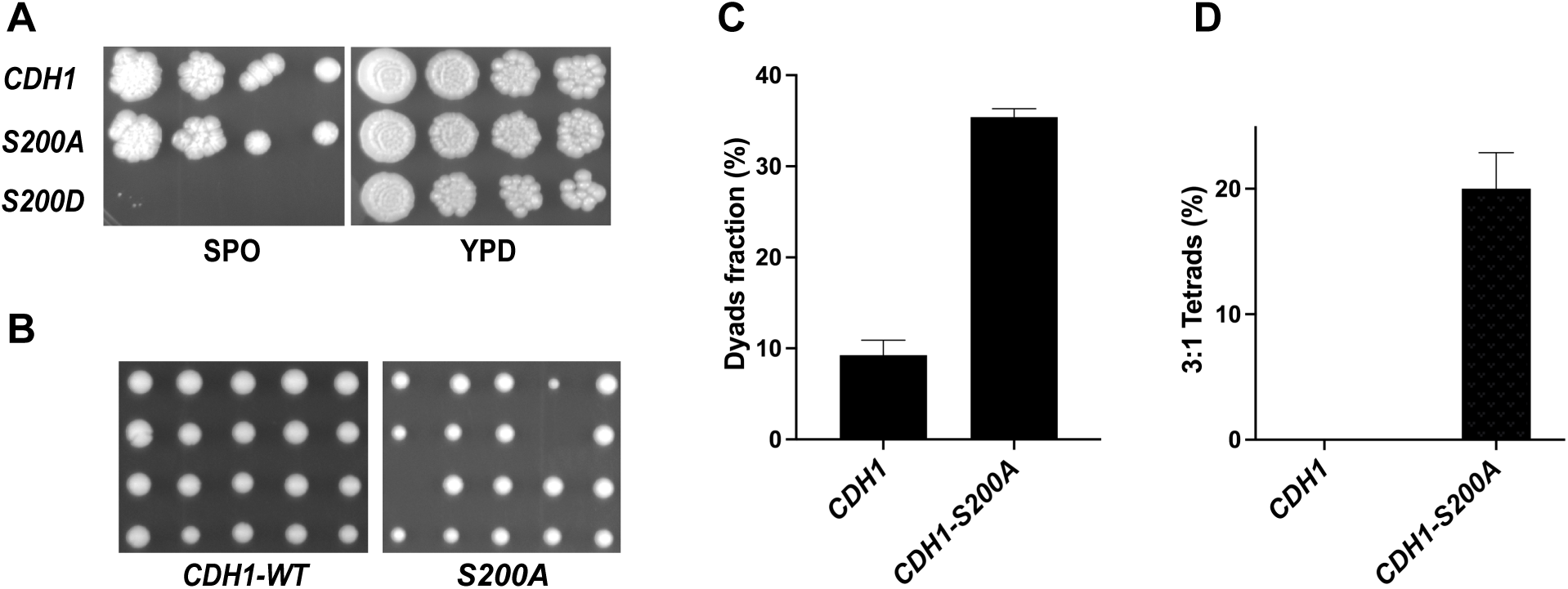
Stabilized Cdh1-S200A increases the rate of chromosome mis-segregation during sporulation. **(A)** Diploid wild-type *CDH1* (strain DOY2899), *CDH1-S200A* (strain DOY4191), and *CDH1-S200D* (strain DO3970) cells were incubated in sporulation-inducing medium (SPO) for 7 days at 30°C, serially diluted, and spotted on YPD plates (*left*). Asynchronous cultures of the same strains grown in rich medium (YPD) to early stationary phase were similarly diluted and spotted (*right*); both plates were grown at 30°C for three days. **(B)** Following incubation in sporulation-inducing medium, tetrads derived from wild type *CDH1* (strain DOY2899) and *CDH1-S200A* (strain DO4191) cells were dissected, and individual spores were spotted on YPD plates. The resulting spores germinated into colonies of haploid cells for 3 days at 30°C. Of note, strain *CDH1-S200D* (strain DOY3970) did not form any tetrads in sporulation-inducing medium and, thus, was impossible to dissect. **(C)** Incomplete meiosis or sporulation leads to the appearance of dyads, composed of only two spores. The fraction of dyads in 500 random tetrads from wild type *CDH1* (strain DOY2899) and isogenic *CDH1-S200A* (strain DO4191) strains is shown. **(D)** Under optimal sporulation conditions, yeast diploid cells produce 4 viable haploid spores. Wild type *CDH1* (strain DOY2899) and isogenic *CDH1-S200A* (strain DO4191) cells were sporulated for 2 days at 30°C, approximately 100 asci were dissected, and the percentage of tetrads with only 3 viable spores (3:1 Tetrads) were plotted. Examples of such tetrads are shown in panel B.

Microscopic examination revealed sporulation defects in *CDH1-S200A* cells. In particular, about four times as many *CDH1-S200A* cells than wild-type cells formed dyads rather than tetrads (Figure 6C), suggesting that these cells completed only one round of cell division following meiotic DNA replication. Moreover, about 20% of the tetrads formed from *CDH1-S200A* cells contained only three viable spores (Figures 6B and 6D), with one spore unable to undergo even a first round of cell division. In contrast, all 100 tetrads formed from wild-type cells produced four viable spores (Figures 6B and 6D).

## DISCUSSION

The activities of key cell cycle regulators, such as APC/C^Cdh1^, in yeast and in higher eukaryotes are tightly regulated through the integration of signals from internal and external sources. We have discovered a complex network of signals leading to the abrupt degradation of Cdh1 during the transition from vegetative growth to the meiotic/sporulation program. Destabilization of Cdh1 required heterozygosity at the mating-type locus (*MATa/MATα* cells) and starvation for glucose. The apparent degradation of Cdh1 was mediated by the 26S proteasome and involved “auto-ubiquitination” of Cdh1 by APC/C^Cdh1^ operating *in trans*. We found that the glucose-sensing branch of this regulatory system runs through the SNF1 protein kinase complex, which directly phosphorylates Cdh1 on Ser-200, leading to its degradation. Failure to degrade Cdh1 leads to chromosome instability and aneuploidy, as well as a high rate of meiosis II failure, leading to formation of dyads rather than tetrads.

There are striking similarities and differences in the regulation of Cdh1 stability by phosphorylation in human and yeast cells. In both species, the Cdh1 N-terminal domain is mostly unstructured and contains multiple sites of phosphorylation regulating protein-protein interactions and protein stability. Several of these sites are evolutionary conserved, as are some of the protein kinases responsible for Cdh1 phosphorylation. The most prominent such protein kinase is CDK, which phosphorylates at least nine distinct sites within the N-terminal region of yeast Cdh1, thereby controlling protein localization and its binding to the core APC/C subunits. In human cells, phosphorylation by cyclin A-CDK primes Cdh1 for subsequent phosphorylation by Plk1, which is followed by β3-TRCP binding and ubiquitination of Cdh1 by the SCF ^β3-TRCP^ ubiquitin ligase (Fukushima *et al*, 2013; Lukas *et al*, 1999; Sorensen *et al*, 2000). In addition, cyclin F, another SCF adaptor protein, targets Cdh1 for SCF-mediated ubiquitination during early S-phase (Choudhury *et al*, 2016). Given the striking conservation of the yeast and human systems, we were surprised by the lack of involvement of SCF complexes in the degradation of Cdh1 during meiosis and sporulation. Rather, degradation appears to be totally dependent on the APC/C.

How the APC/C targets Cdh1 for degradation was surprising and quite different from how it recognizes typical APC/C substrates. First, yeast Cdh1 does not contain any classical D-box, KEN box or ABBA motifs. Deletion analysis failed to identify any small region necessary for degradation. Second, we found that Cdh1 degradation occurred in *trans-*, i.e., an APC/C^Cdh1^ complex recognizes a second, substrate molecule, of Cdh1 for ubiquitination. This mode of recognition contrasts with the “auto-ubiquitination” of Cdc20 within the APC/C^Cdc20^ complex during mitosis (Foe *et al*., 2011). Third, Cdh1 degradation required a priming phosphorylation on Ser-200. Phosphorylation-dependent regulation has been previously reported for several APC/C substrates, including FBXO31, Mps1, Fin1, Pds1 and Nrm1 (Choppara *et al*, 2018) (Holt *et al*, 2008; Jaspersen *et al*, 2004; Ostapenko & Solomon, 2011; Woodbury & Morgan, 2007).

It is interesting that the degradation of Cdh1, which is necessary for proper meiosis and sporulation, doesn’t actually require that cells initiate the meiotic program. Since Cdh1 degradation occurs even in *ime1Δ* cells, it seems that the mating type and nutritional signals that tell cells to enter the meiotic program independently signal the degradation of Cdh1. Thus, in searching for proteins involved in the decision to degrade Cdh1, we can at once exclude all early, middle, and late meiosis-specific gene products. Since cells enter meiosis upon glucose and nitrogen limitation, it was not entirely surprising that SNF1 activity was required for Cdh1 degradation. In fact, the SNF1 protein kinase complex acts independently of Ime1, and so is well positioned to be part of a signaling pathway leading to Cdh1 instability.

Our findings lead to additional questions. For instance, how does one molecule of Cdh1 direct another to the APC/C for ubiquitination? Phosphorylation of Ser-200 may create a motif recognized directly by the second Cdh1 molecule, or it may induce a conformational change in another region of Cdh1, allowing its recognition. As noted earlier (see Supplemental Figure 3), sequences between amino acids 50 and approximately 150 appear to play a role in Cdh1 degradation, and could include this novel degron. A second question is what protein normally expressed in diploids is necessary, along with glucose starvation, to induce the degradation of Cdh1. As we showed, diploidy *per se* is not necessary, as Cdh1 is degraded in *MATa/MATα* haploid cells, so the “diploid factor” is downstream of the a1-alpha2 transcription factor. There are currently no obvious candidates for this factor. Finally, we presume that Cdh1 must be degraded during meiosis so that an APC/C substrate important for proper spindle assembly or chromosomal segregation can persist until the end of meiosis. The key feature of meiosis—a reductional division followed immediately by an equational division without intervening Gap or S phases—requires that many proteins that would normally be degraded at the end of MI persist into MII. Such proteins include cyclins (Clb1 and Clb3) as well as regulators of chromosome cohesion, Cdc5 (polo-like kinase), Cdc20, Mps1, Pds1, and Sgo1. Three APC/C activators— Cdh1, Cdc20, and Ama1—are each essential for meiosis and sporulation. The timing of synthesis, degradation, and activity of each sets the choreography of the complex cell cycle program of meiosis. These findings may lead to novel approaches in the development of anti-fungal medications. Although Ser-200 is not conserved in mammalian Cdh1, the general scheme outlined here may nevertheless have implications for understanding the development of aneuploidy in human development, which can lead to Down and Edwards Syndromes (trisomy 21 and 18, respectively), trisomy for the sex chromosomes, as well as fetal inviability.

## ACKNOWLEDGEMENTS

The work reported here is substantially revised from a previous submission to bioRxiv (doi: https://doi.org/10.1101/358655). This work was supported by grant R01 GM129155 (to M.J.S.) from the NIH.

## MATERIALS AND METHODS

### Yeast strains and plasmids

Yeast strains were derivatives of SK1 (*ho::LYS2 leu2::hisG his3::hisG trp::hisG ura3::pGPD1-GAL4(848).ER::URA3*) and W303 (*ade2-1 trp1-1 leu2-3,112 his3-11,15 ura3-1*); their relevant genotypes are listed in Supplementary Table 1. The SK1 strains were provided by Angelika Amon (MIT, Cambridge, MA) (Carlile & Amon, 2008). Conditional *cdc23-1* and *cdc34-2* strains were described previously (Burton & Solomon, 2000). The *MATa/MATa* diploid strain and *MATalpha2* expression vector were provided by Mark Hochstrasser (Yale University, New Haven, CT) (Hickey & Hochstrasser, 2015). The *tpk1-as tpk2Δ tpk3Δ* strain was a generous gift of Angelika Amon (MIT, Cambridge, MA) (Weidberg *et al*, 2016). The *cdh1-dbr, cdh1-kbr, cdh1-amr* strains were provided by Mark Hall (Purdue University, West Lafayette, IN) (Qin *et al*., 2016). The *Myc_9_-CDH1* plasmid was provided by Wolfgang Seufert (University of Stuttgart, Stuttgart, Germany) (Zachariae *et al*., 1998) (Zachariae, Schwab et al. 1998). *Myc_9_-CDH1* construction of *ime1Δ* (W303 *ime1::natMX4*), *ime2Δ* (W303 *ime2::natMX4*), *snf1Δ* (W303 *snf1::natMX4*), *snf4Δ* (W303 *snf4::natMX4*), *sip1Δ* (W303 *sip1::natMX4*), *sip2Δ* (W303 *sip2::natMX4*), *gal83Δ* (W303 *gal83::natMX4*), *elm1Δ* (W303 *elm1::natMX4*), *mig1Δ* (W303 *mig1::natMX4*), *adr1Δ* (W303 *adr1::natMX4*), *tor1Δ* (W303a *tor1::natMX4*), and *sak1Δ* (W303a *sak1:: natMX4*) strains were accomplished by a PCR-based method (Goldstein & McCusker, 1999). Gene disruptions were verified by PCR using a primer downstream of the deleted gene and a primer internal to *natMX4*.

The *Myc_9_-CDH1* expression plasmid was used as template to introduce alanine or aspartic acid mutations within phospho-acceptor sites: S200, S418 and a cluster of CDK phospho-acceptor sites: S157, S169, T173, T176, S227, S239. The following in-frame deletions were introduced: *CDH1-Δ1-50, CDH1-Δ1-80, CDH1-Δ1-150, CDH1-Δ50-157.* The *GST-CDH1* expression plasmid was described previously (Burton & Solomon, 2000). The *GST-SNF1-G53R* expression vector was constructed by cloning *SNF1-G53R* (Orlova *et al*, 2006) between *BamH*I and *Sal*I sites of pEG-KT. All mutations were verified by sequencing of the entire coding region. Primer sequences and further details of the plasmids are available upon request.

### Cell growth and sporulation conditions

Cultures were grown in YPD, YPA (1% yeast extract, 2% peptone, 2% potassium acetate), and in complete minimal (CM) media as described (Guthrie & Fink, 1991). For sporulation time courses, cells were grown overnight in YPD, diluted into YPA or CM-Raffinose (OD_600_ ∼0.3) and grown for 12 hours at 30°C. Cells were washed and released into sporulation medium (SPM) containing 0.3% potassium acetate, 0.02% raffinose, 20 µg/ml mixture of essential amino acids, adenine, and uracil for 24 hours at 30°C. Alternatively, conditional mutant *cdc23-1* and *cdc34-2* cells were grown at 23°C in CM-Raffinose and released into SPM at 35°C for 24 hours. Starting from the release, equal volumes of cells (8 ml) were collected at 6-hour intervals by centrifugation at 4000 rpm for 2 min and pellets were frozen in liquid nitrogen. For analysis of APC/C^Cdh1^ substrates, haploid *MATa bar1Δ* cells were grown in YPD to mid-exponential phase (OD_600_ ∼0.5), and arrested in G1 phase in the presence of 100 ng/ml *a*-factor for 2 hours at 30°C.

For tetrad analysis and dissection, cells were transferred from CM-Raffinose (OD_600_ ∼0.3) to sporulation medium (SPM) and incubated for 3 days at 30°C. Sporulating cells were harvested and incubated with 0.5 mg/ml Zymolyase-100T (ICN Immunobiologicals) in 1M sorbitol for 10 min at 30°C, washed, and microscopically dissected on YPD plates. The resulting spores were grown for 3 days at 30°C.

### Yeast extracts and Immunoblotting

Cells from 8 ml cultures (OD_600_ ∼0.4) were collected, washed with ice-cold H_2_O, suspended in 0.5 ml 100 mM NaOH and incubated for 5 min at 23°C. Cells were pelleted, suspended in 0.1 ml of 1xSDS loading buffer (60 mM Tris-Cl pH 6.8, 2% SDS, 5% glycerol, 100 mM β-ME) and lysed by heating at 95°C for 5 min. Cell debris was removed by centrifugation at 14,000 rpm in a microfuge at 23°C for 5 min. Yeast protein extracts were separated on protein gels containing 10% polyacrylamide and transferred to an Immobilon-P membrane (Millipore). The membranes were probed with anti-Myc antibodies (9E10, 1 µg/ml, Millipore) overnight in Blotto (10 mM Tris-Cl pH 7.5, 150 mM NaCl, 0.1% Tween-20, 5% dry milk) at 4°C. Membranes were then washed with TBST and incubated with anti-mouse horseradish peroxidase (HRP)-conjugated secondary antibodies (1:1000, Pierce). Proteins were visualized by chemiluminescence (SuperSignal, Pierce).

### Protein kinase assays

Recombinant Cdh1^1-250^-His_6_ was expressed from the pGEX-6M vector in *E. coli* and purified on Talon resin as previously described (Ostapenko *et al*, 2008). The SNF1 protein kinase was purified from strain DOY4195 (*MATa pep4-3 his3-1 leu2-3112 pEG-KT-SNF1-G53R*) yeast using a single-step affinity purification. Briefly, 1 l of DOY4195 cells were grown in YP-Raffinose to OD_600_ = 0.5, induced with 2% galactose for 5 hours at 30°C, washed, and pelleted by centrifugation at 5,000 rpm for 10 min at 4°C. Cells were lysed by shaking 5 ml suspensions with glass beads (0.5-mm diameter; Sigma) in a bead beater (Biospec Products) in 50 mM Hepes, pH 7.5, 150 mM NaCl, 5 mM EDTA, 10% glycerol, 1 mM DTT, 0.5 mM PMSF, 1 tablet of protease inhibitors (cOmplete EDTA-free, Roche). The cell extract was clarified by centrifugation at 30,000 rpm in a TLA.103 rotor (Beckman, Brea, CA) for 20 min at 4°C; GST-SNF1 was purified using 0.5 ml bed volume of Glutathione-Sepharose resin (GE Healthcare), aliquoted, and stored at-80°C.

For kinase assays, approximately 1.0 µg GST-SNF1 on glutathione beads was mixed with 1.0 µg Cdh1^1-250^-His_6_ in 50 mM Hepes, pH 7.5, 150 mM NaCl, 30 mM MgCl_2_, 0.5 mM DTT, 1 mM ATP in a 50-µl volume and incubated for 30 min at 23°C. The reaction products were separated on a Phos-tag^TM^ (Fujifilm WAKO, Japan) polyacrylamide gel in the presence of 10 mM MnCl_2_, according to manufacturer’s instructions. Proteins were detected by immunoblotting with anti-Cdh1 antibodies (1 µg/ml, Millipore) and anti-GST antibodies (0.5 µg/ml, Invitrogen). Membranes were then incubated with anti-rabbit HRP-conjugated secondary antibodies (1:1000, Pierce) and visualized by chemiluminescence (SuperSignal, Pierce).

---

**Supplementary Table 1.**
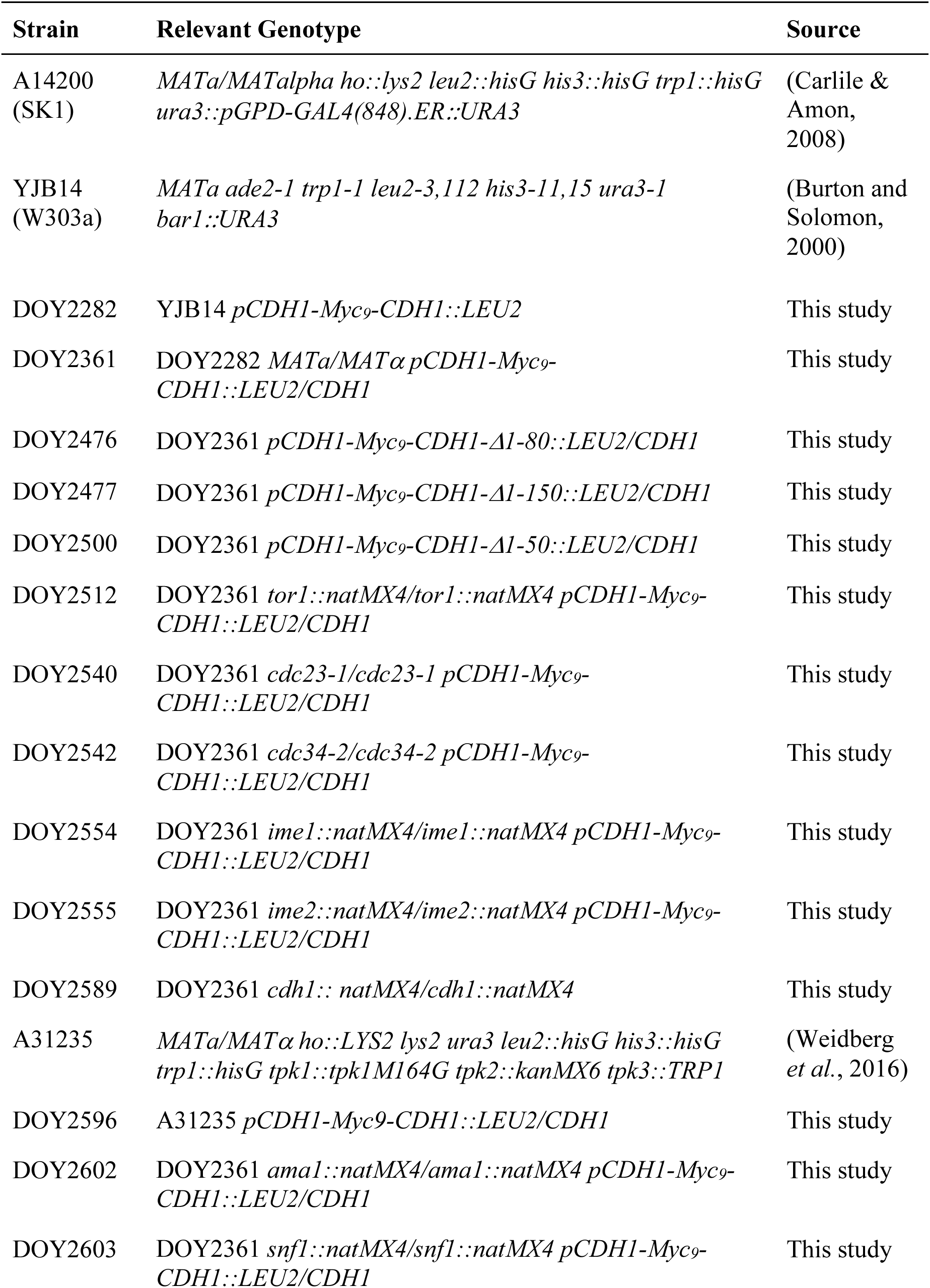
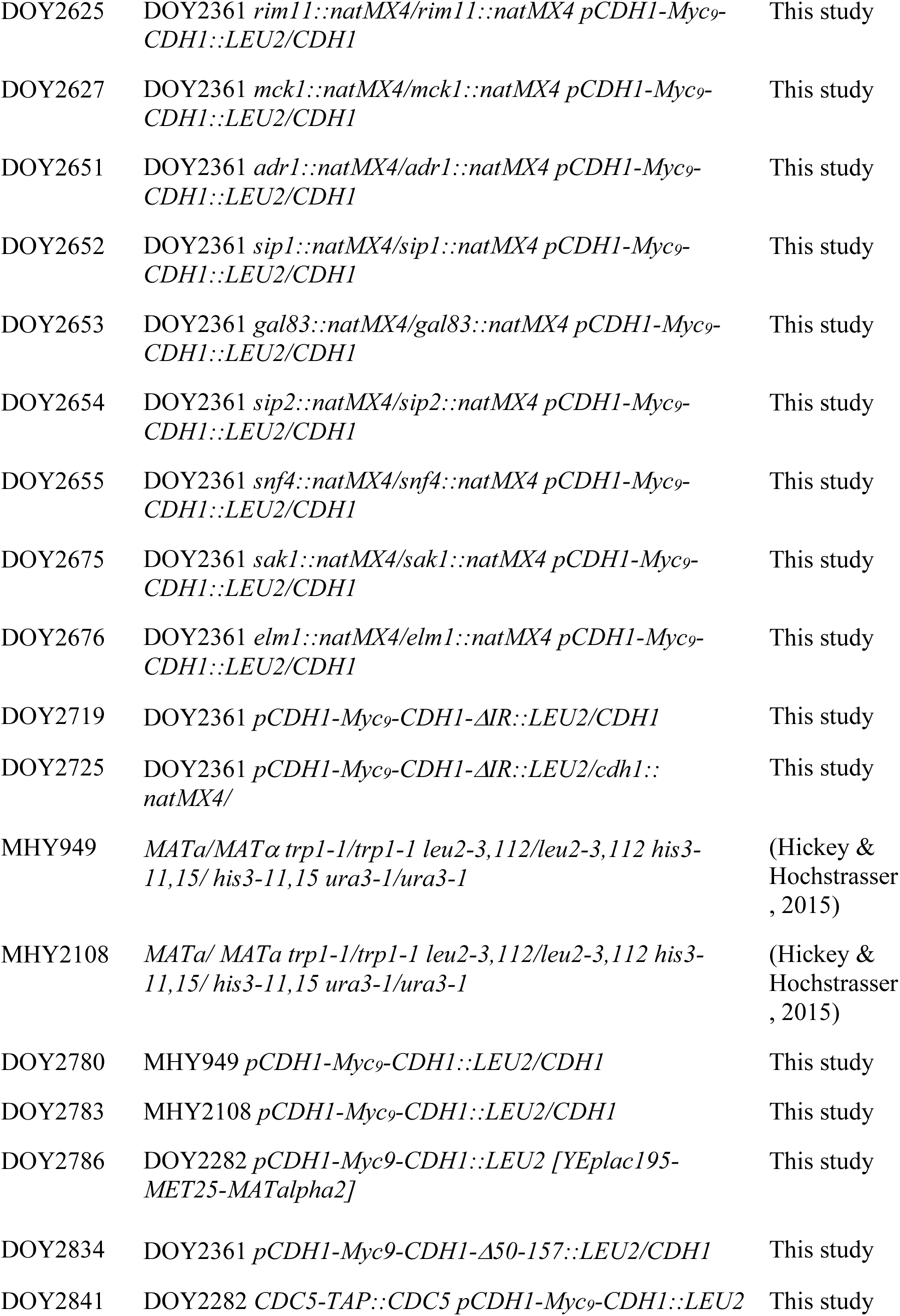
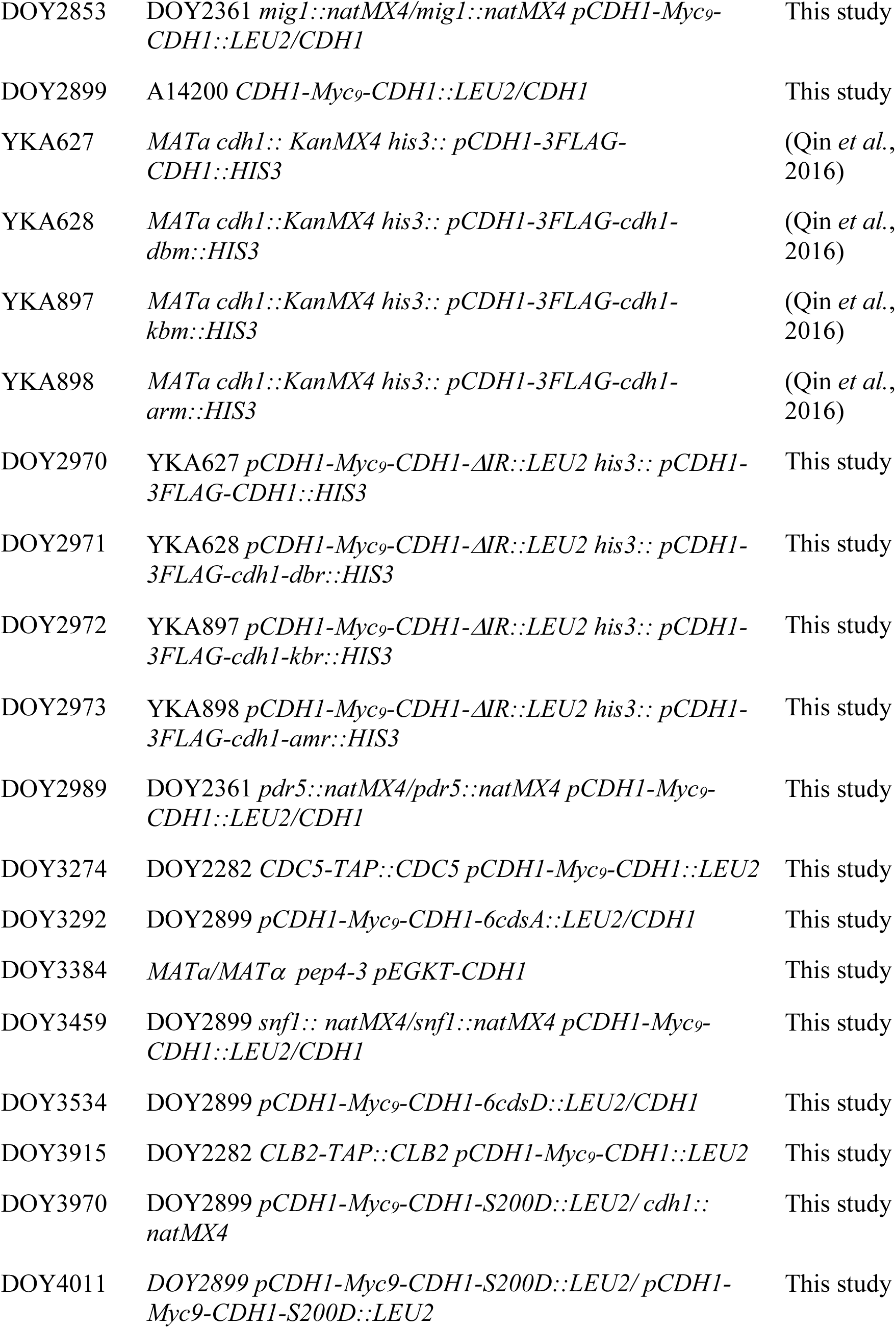
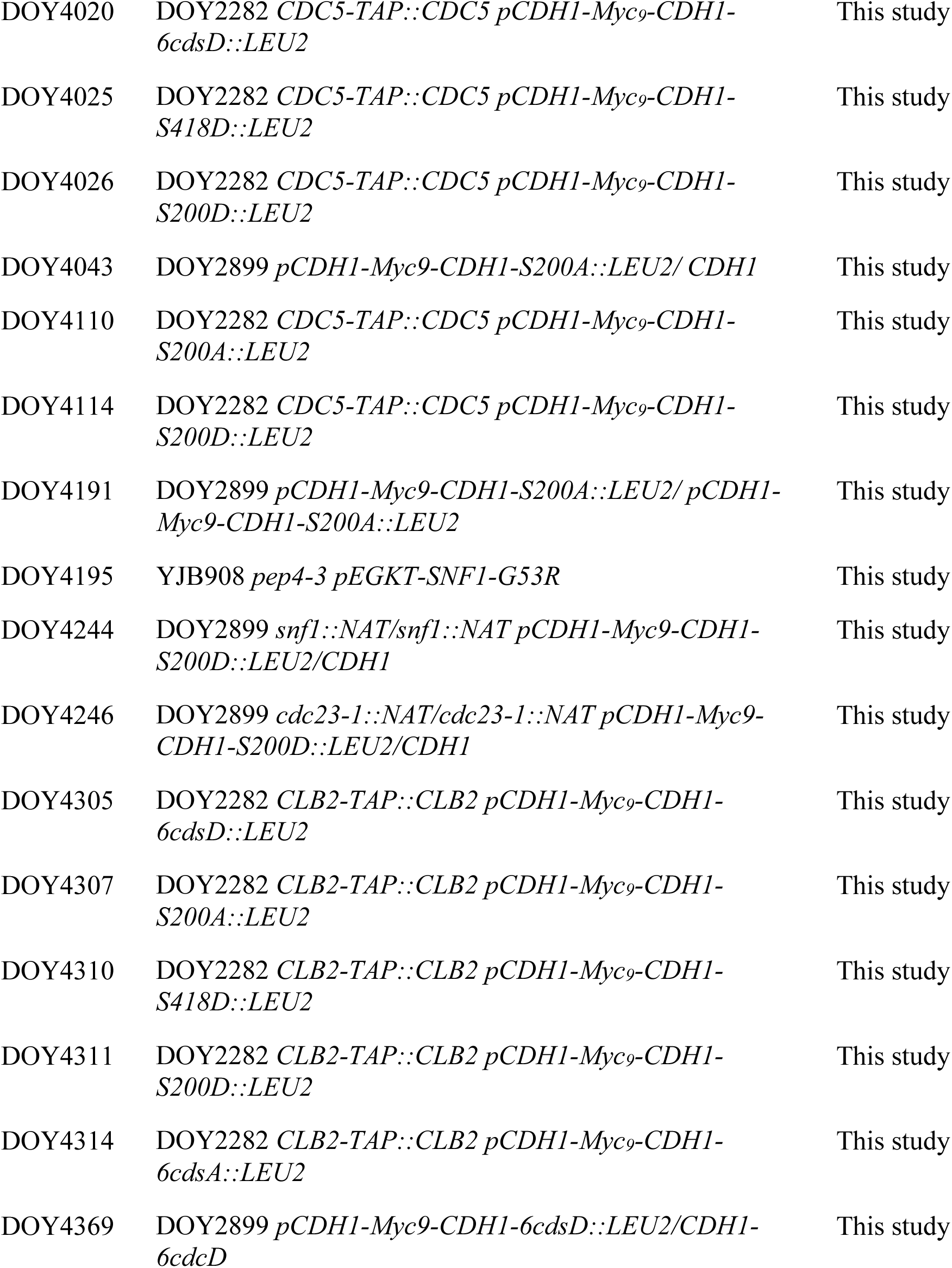
*S. cerevisiae* strains used in this study.

**Figure 3 Supplementary.**
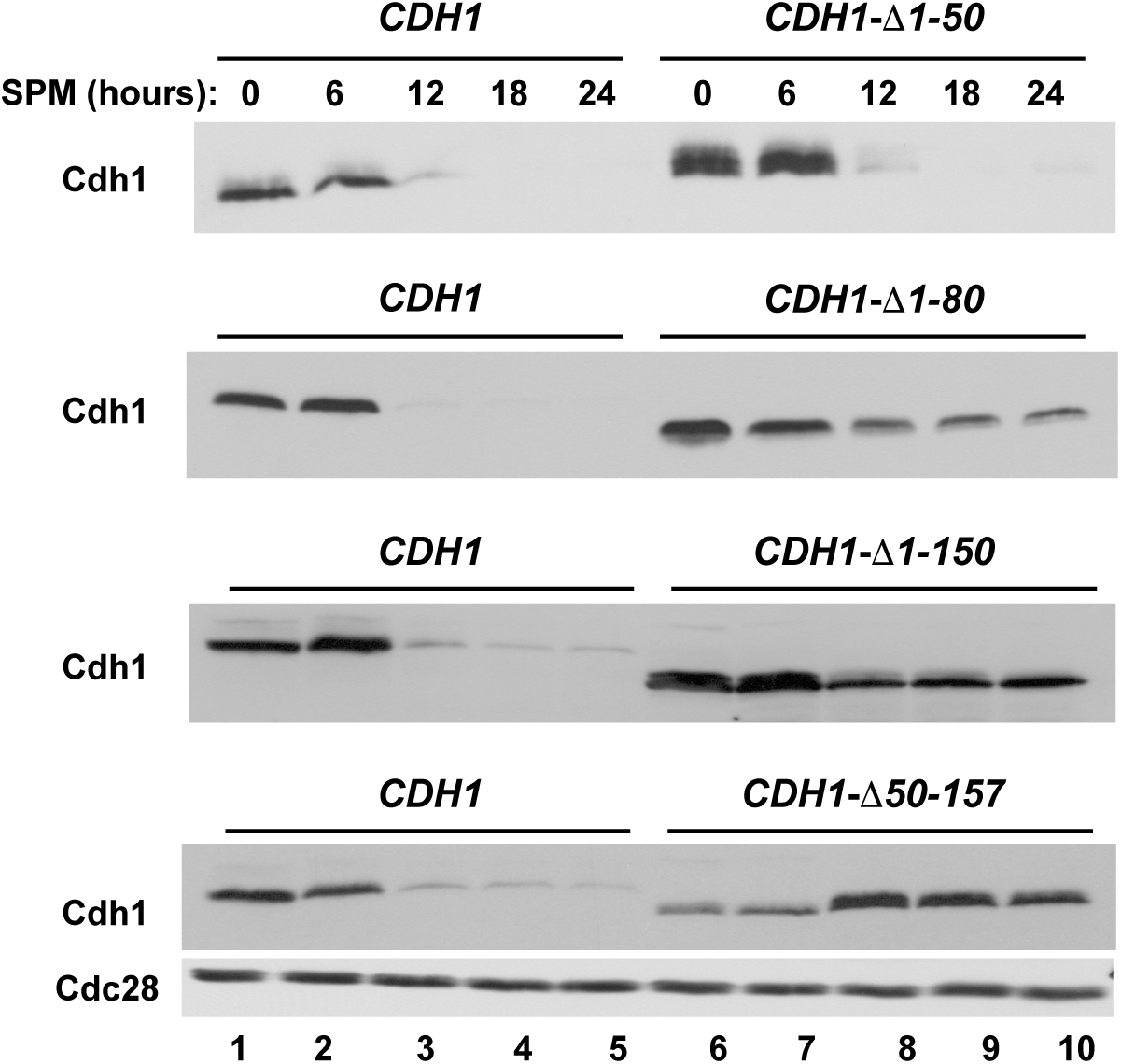
N-terminal sequences are necessary for Cdh1 degradation. Endogenous wild-type *Myc-CDH1* and the indicated truncations were co-expressed with an untagged copy of *CDH1* in *MATa/MATα* diploid strains. The resulting heterozygous strains (*Myc-CDH1*/*CDH1* (strain DOY2361), *Myc-CDH1-Δ1-50*/*CDH1* (strain DOY2500)*, Myc-CDH1-Δ1-80*/*CDH1* (strain DOY2476)*, Myc-CDH1-Δ1-150*/*CDH1* (strain DOY2477)*, and Myc-CDH1-Δ50-157*/*CDH1)* (strain DOY2834) were tested for Myc-Cdh1 stability after transition to sporulation medium. Yeast extracts prepared from the samples withdrawn at the indicated times were immunoblotted with anti-Myc antibodies to detect Cdh1. The membranes were then re-probed with anti-Cdc28 antibodies to serve as a loading control.

**Figure 4 Supplementary.**
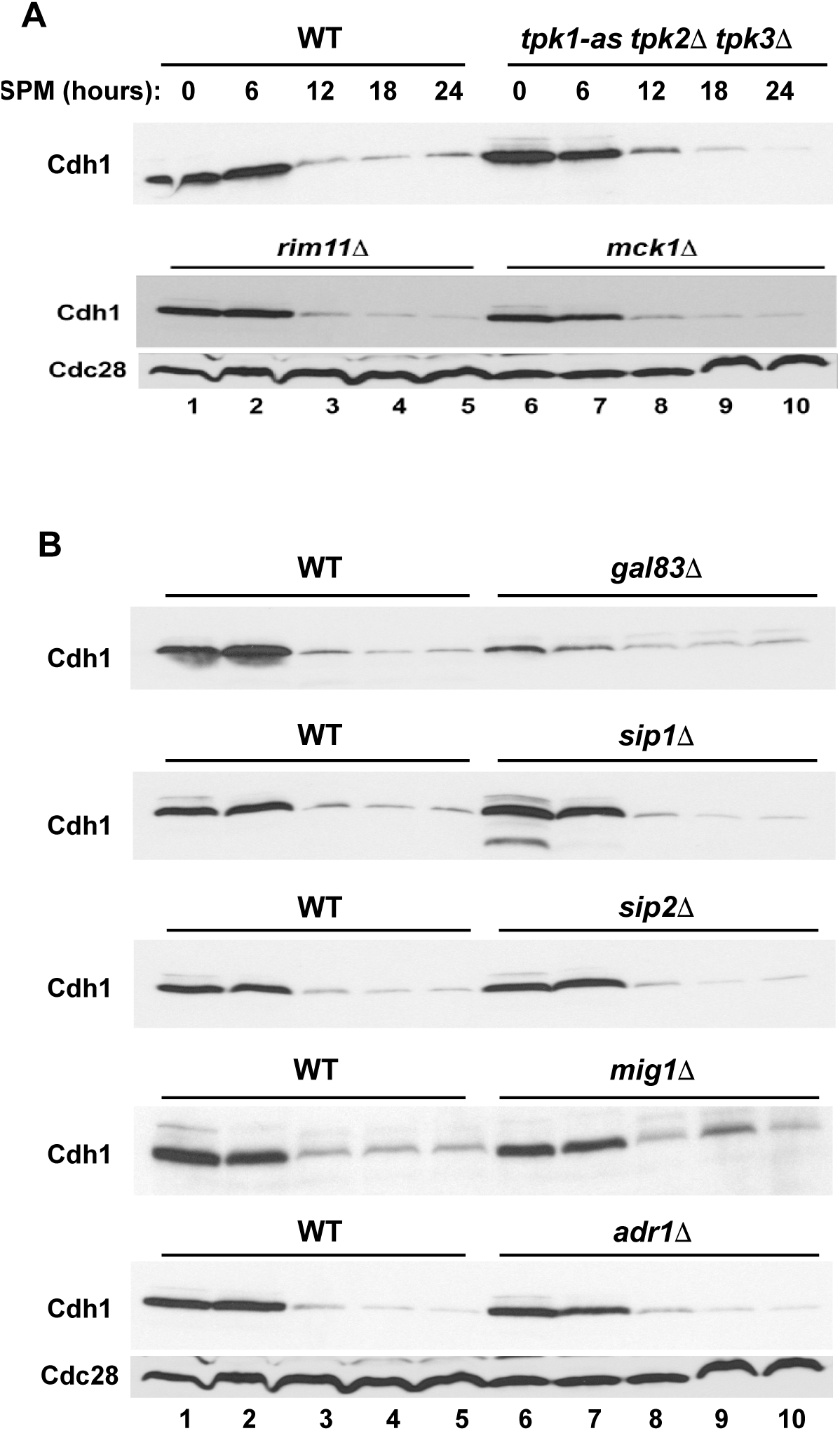
Redundant subunits of SNF1 and downstream transcription factors are not required for Cdh1 turnover. **(A)** (*top*) cAMP-dependent protein kinase A (PKA) activity can be eliminated by deleting two of the three PKA catalytic subunit isoforms (Tpk2, Tpk3) and expressing an analog-sensitive version of the third isoform (Tpk1, *tpk1-as*). Homozygous cells (strain DOY2596) carrying *tpk1-as tpk2Δ tpk3Δ* mutations were grown in YPA, transferred to sporulation medium in the absence (lanes 1-5) or presence or 1 µg/ml of 1NM-PP1 to rapidly inactivate Tpk1-as (lanes 6-10). (*bottom*) Wild-type (strain DOY2361) and isogenic diploid yeast strains carrying deletions of meiosis-specific kinases *RIM11* (strain DOY2625) and *MCK1* (strain DOY2627), were grown in YPD and transferred to sporulation-inducing medium as in Figure 1A. Samples were withdrawn at the indicated times and probed to measure Cdh1 protein levels with 9E10 antibodies. The membranes were then re-probed with anti-PSTAIR antibodies to detect Cdc28 as a loading control. **(B)** (*top three panels*) Wild-type (strain DOY2361) and isogenic diploid yeast strains carrying deletions of SNF1 β-subunits, *GAL83* (strain DOY2653), *SIP1* (strain DOY2652), *SIP2* (strain DOY2654)-were grown in YPA and transferred to sporulation medium. (*bottom two panels*) A sporulation time course was carried with yeast strains carrying mutations of transcription factors *MIG1* (*mig1Δ* strain DOY2853) and *ADR1* (*adr1Δ* strain DOY2651) that function downstream of Snf1 pathway. Wild type and the indicated mutant strains were grown in YPA, transferred to sporulation medium for indicated time intervals. Samples were processed for immunoblotting to detect Myc-Cdh1 as in Figure 1A. Anti-Cdc28 probing was used as a loading control.

**Figure 5 Supplementary.**
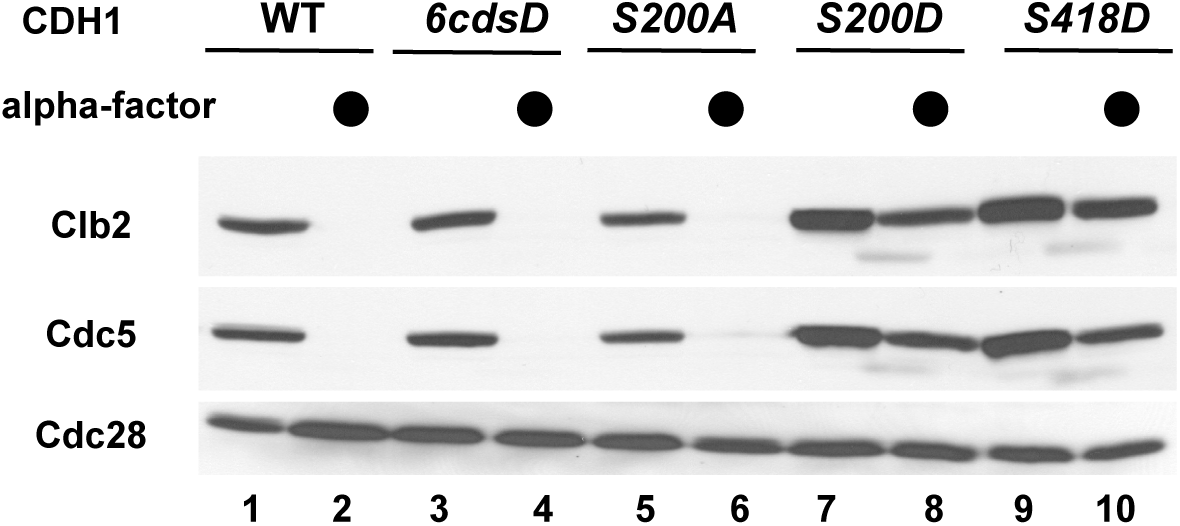
Cdh1-S200A can promote APC/C substrate degradation. Isogenic haploid *MATa* strains carrying endogenous *CDH1* (strains DOY3915 and DOY2841), *CDH1-6cdsD* (strains DOY4305 and DOY4020), *CDH1-S200A* (strains DOY4307 and DOY4110), *CDH1-S200D* (strains DOY4311 and DOY4114), and *CDH1-S418D* (strains DOY4310 and DOY4025) were incubated in the absence (odd lanes) or presence (even lanes) of the mating pheromone alpha-factor for 2 hours at 30°C. Alpha-factor treatment arrests haploid cells in G1, leading to activation of APC/C^Cdh1^ and degradation of its substrates, Clb2 (strains DOY3915, DOY4305, DOY4307, DOY4311, DOY4310) and Clb5 (strains DOY2841, DOY4020, DOY4110, DOY4114, DOY4025). Yeast protein extracts were processed for immunoblotting to detect Clb2-TAP and Cdc5-TAP with PAP antibodies. Immunoblotting for Cdc28 was used as a loading control.

## REFERENCES

Barford D (2011) Structure, function and mechanism of the anaphase promoting complex (APC/C). Q Rev Biophys 44: 153–190

Brandeis M, Hunt T (1996) The proteolysis of mitotic cyclins in mammalian cells persists from the end of mitosis until the onset of S phase. EMBO J 15: 5280–5289

Brown NG, VanderLinden R, Watson ER, Qiao R, Grace CR, Yamaguchi M, Weissmann F, Frye JJ, Dube P, Ei Cho S et al. (2015) RING E3 mechanism for ubiquitin ligation to a disordered substrate visualized for human anaphase-promoting complex. Proc Natl Acad Sci U S A 112: 5272–5279

Burton JL, Solomon MJ (2000) Hsl1p, a Swe1p inhibitor, is degraded via the anaphase-promoting complex. Mol Cell Biol 20: 4614–4625

Cardozo T, Pagano M (2004) The SCF ubiquitin ligase: insights into a molecular machine. Nature Reviews Molecular Cell Biology 5: 739–751

Carlile TM, Amon A (2008) Meiosis I is established through division-specific translational control of a cyclin. Cell 133: 280–291

Chang L, Zhang Z, Yang J, McLaughlin SH, Barford D (2015) Atomic structure of the APC/C and its mechanism of protein ubiquitination. Nature 522: 450–454

Choppara S, Malonia SK, Sankaran G, Green MR, Santra MK (2018) Degradation of FBXO31 by APC/C is regulated by AKT-and ATM-mediated phosphorylation. Proc Natl Acad Sci U S A 115: 998–1003

Choudhury R, Bonacci T, Arceci A, Lahiri D, Mills CA, Kernan JL, Branigan TB, DeCaprio JA, Burke DJ, Emanuele MJ (2016) APC/C and SCF(cyclin F) Constitute a Reciprocal Feedback Circuit Controlling S-Phase Entry. Cell Rep 16: 3359–3372

Cooper KF, Mallory MJ, Egeland DB, Jarnik M, Strich R (2000) Ama1p is a meiosis-specific regulator of the anaphase promoting complex/cyclosome in yeast. Proc Natl Acad Sci U S A 97: 14548–14553

Cooper KF, Strich R (2011) Meiotic control of the APC/C: similarities & differences from mitosis. Cell Div 6: 16

Crasta K, Lim HH, Giddings TH, Jr., Winey M, Surana U (2008) Inactivation of Cdh1 by synergistic action of Cdk1 and polo kinase is necessary for proper assembly of the mitotic spindle. Nat Cell Biol 10: 665–675

Davey NE, Morgan DO (2016) Building a Regulatory Network with Short Linear Sequence Motifs: Lessons from the Degrons of the Anaphase-Promoting Complex. Mol Cell 64: 12–23

Di Fiore B, Davey NE, Hagting A, Izawa D, Mansfeld J, Gibson TJ, Pines J (2015) The ABBA motif binds APC/C activators and is shared by APC/C substrates and regulators. Dev Cell 32: 358–372

Estruch F, Treitel MA, Yang X, Carlson M (1992) N-terminal mutations modulate yeast SNF1 protein kinase function. Genetics 132: 639–650

Foe IT, Foster SA, Cheung SK, DeLuca SZ, Morgan DO, Toczyski DP (2011) Ubiquitination of Cdc20 by the APC occurs through an intramolecular mechanism. Curr Biol 21: 1870–1877

Foster SA, Morgan DO (2012) The APC/C subunit Mnd2/Apc15 promotes Cdc20 autoubiquitination and spindle assembly checkpoint inactivation. Mol Cell 47: 921–932

Fukushima H, Ogura K, Wan L, Lu Y, Li V, Gao D, Liu P, Lau AW, Wu T, Kirschner MW et al (2013) SCF-mediated Cdh1 degradation defines a negative feedback system that coordinates cell-cycle progression. Cell Rep 4: 803–816

Glotzer M, Murray AW, Kirschner MW (1991) Cyclin is degraded by the ubiquitin pathway. Nature 349: 132–138

Goldstein AL, McCusker JH (1999) Three new dominant drug resistance cassettes for gene disruption in Saccharomyces cerevisiae. Yeast 15: 1541–1553

Guthrie C, Fink GR (1991) Guide to yeast genetics and molecular biology. Academic Press, San Diego, CA

He J, Chao WC, Zhang Z, Yang J, Cronin N, Barford D (2013) Insights into degron recognition by APC/C coactivators from the structure of an Acm1-Cdh1 complex. Mol Cell 50: 649–660

Hickey CM, Hochstrasser M (2015) STUbL-mediated degradation of the transcription factor MATalpha2 requires degradation elements that coincide with corepressor binding sites. Mol Biol Cell 26: 3401–3412

Hockner S, Neumann-Arnold L, Seufert W (2016) Dual control by Cdk1 phosphorylation of the budding yeast APC/C ubiquitin ligase activator Cdh1. Mol Biol Cell 27: 2198–2212

Holt LJ, Hutti JE, Cantley LC, Morgan DO (2007) Evolution of Ime2 phosphorylation sites on Cdk1 substrates provides a mechanism to limit the effects of the phosphatase Cdc14 in meiosis. Mol Cell 25: 689–702

Holt LJ, Krutchinsky AN, Morgan DO (2008) Positive feedback sharpens the anaphase switch. Nature 454: 353–357

Irniger S, Nasmyth K (1997) The anaphase-promoting complex is required in G1 arrested yeast cells to inhibit B-type cyclin accumulation and to prevent uncontrolled entry into S-phase. J Cell Sci 110: 1523–1531

Jaquenoud M, F. vD, Peter M (2002) Cell cycle-dependent nuclear export of Cdh1p may contribute to the inactivation of APC/C^Cdh1^. EMBO J 21: 6515–6526

Jaspersen SL, Charles JF, Morgan DO (1999) Inhibitory phosphorylation of the APC regulator Hct1 is controlled by the kinase Cdc28 and the phosphatase Cdc14. Curr Biol 9: 227–236

Jaspersen SL, Huneycutt BJ, Giddings TH, Jr., Resing KA, Ahn NG, Winey M (2004) Cdc28/Cdk1 regulates spindle pole body duplication through phosphorylation of Spc42 and Mps1. Dev Cell 7: 263–274

Kassir Y, Granot D, Simchen G (1988) IME1, a positive regulator gene of meiosis in S. cerevisiae. Cell 52: 853–862

Kinoshita E, Kinoshita-Kikuta E, Takiyama K, Koike T (2006) Phosphate-binding tag, a new tool to visualize phosphorylated proteins. Mol Cell Proteomics 5: 749–757

Lukas C, Sorensen CS, Kramer E, Santoni-Rugiu E, Lindeneg C, Peters JM, Bartek J, Lukas J (1999) Accumulation of cyclin B1 requires E2F and cyclin-A-dependent rearrangement of the anaphase-promoting complex. Nature 401: 815–818

Mallory MJ, Cooper KF, Strich R (2007) Meiosis-specific destruction of the Ume6p repressor by the Cdc20-directed APC/C. Mol Cell 27: 951–961

Neiman AM (2011) Sporulation in the budding yeast Saccharomyces cerevisiae. Genetics 189: 737–765

Okaz E, Arguello-Miranda O, Bogdanova A, Vinod PK, Lipp JJ, Markova Z, Zagoriy I, Novak B, Zachariae W (2012) Meiotic prophase requires proteolysis of M phase regulators mediated by the meiosis-specific APC/CAma1. Cell 151: 603–618

Orlova M, Kanter E, Krakovich D, Kuchin S (2006) Nitrogen availability and TOR regulate the Snf1 protein kinase in Saccharomyces cerevisiae. Eukaryot Cell 5: 1831–1837

Ostapenko D, Burton JL, Wang R, Solomon MJ (2008) Pseudosubstrate inhibition of the anaphase-promoting complex by Acm1: regulation by proteolysis and Cdc28 phosphorylation. Mol Cell Biol 28: 4653–4664

Ostapenko D, Solomon MJ (2011) Anaphase promoting complex-dependent degradation of transcriptional repressors Nrm1 and Yhp1 in Saccharomyces cerevisiae. Mol Biol Cell 22: 2175–2184

Penkner AM, Prinz S, Ferscha S, Klein F (2005) Mnd2, an essential antagonist of the anaphase-promoting complex during meiotic prophase. Cell 120: 789–801

Peters JM (2006) The anaphase promoting complex/cyclosome: a machine designed to destroy. Nature Reviews Molecular Cell Biology 7: 644–656

Pfleger CM, Lee E, Kirschner MW (2001) Substrate recognition by the Cdc20 and Cdh1 components of the anaphase-promoting complex. Genes Dev 15: 2396–2407.

Pines J (2011) Cubism and the cell cycle: the many faces of the APC/C. Nat Rev Mol Cell Biol 12: 427–438

Primorac I, Musacchio A (2013) Panta rhei: the APC/C at steady state. J Cell Biol 201: 177–189

Prinz S, Hwang ES, Visintin R, Amon A (1998) The regulation of Cdc20 proteolysis reveals a role for APC components Cdc23 and Cdc27 during S phase and early mitosis. Curr Biol 8: 750–760

Qin L, Guimaraes DS, Melesse M, Hall MC (2016) Substrate Recognition by the Cdh1 Destruction Box Receptor Is a General Requirement for APC/CCdh1-mediated Proteolysis. J Biol Chem 291: 15564–15574

Robbins JA, Cross FR (2010) Requirements and reasons for effective inhibition of the anaphase promoting complex activator CDH1. Mol Biol Cell 21: 914–925

Schwab M, Lutum AS, Seufert W (1997) Yeast Hct1 is a regulator of Clb2 cyclin proteolysis. Cell 90: 683–693

Shirayama M, Zachariae W, Ciosk R, Nasmyth K (1998) The Polo-like kinase Cdc5p and the WD-repeat protein Cdc20p/fizzy are regulators and substrates of the anaphase promoting complex in *Saccharomyces cerevisiae*. EMBO J 17: 1336–1349

Sorensen CS, Lukas C, Kramer ER, Peters JM, Bartek J, Lukas J (2000) Nonperiodic activity of the human anaphase-promoting complex-Cdh1 ubiquitin ligase results in continuous DNA synthesis uncoupled from mitosis. Mol Cell Biol 20: 7613–7623.

Tan GS, Lewandowski R, Mallory MJ, Strich R, Cooper KF (2013) Mutually dependent degradation of Ama1p and Cdc20p terminates APC/C ubiquitin ligase activity at the completion of meiotic development in yeast. Cell Div 8: 9

Tan GS, Magurno J, Cooper KF (2011) Ama1p-activated anaphase-promoting complex regulates the destruction of Cdc20p during meiosis II. Mol Biol Cell 22: 315–326

Thornton BR, Toczyski DP (2006) Precise destruction: an emerging picture of the APC. Genes & Development 20: 3069–3078

Treitel MA, Kuchin S, Carlson M (1998) Snf1 protein kinase regulates phosphorylation of the Mig1 repressor in Saccharomyces cerevisiae. Mol Cell Biol 18: 6273–6280

Visintin C, Tomson BN, Rahal R, Paulson J, Cohen M, Taunton J, Amon A, Visintin R (2008) APC/C-Cdh1-mediated degradation of the Polo kinase Cdc5 promotes the return of Cdc14 into the nucleolus. Genes Dev 22: 79–90

Visintin R, Prinz S, Amon A (1997) *CDC20* and *CDH1*: a family of substrate-specific activators of APC-dependent proteolysis. Science 278: 460–463

Wasch R, Cross FR (2002) APC-dependent proteolysis of the mitotic cyclin Clb2 is essential for mitotic exit. Nature 418: 556–562

Weidberg H, Moretto F, Spedale G, Amon A, van Werven FJ (2016) Nutrient Control of Yeast Gametogenesis Is Mediated by TORC1, PKA and Energy Availability. PLoS Genet 12: e1006075

Woodbury EL, Morgan DO (2007) Cdk and APC activities limit the spindle-stabilizing function of Fin1 to anaphase. Nat Cell Biol 9: 106–112

Zachariae W, Schwab M, Nasmyth K, Seufert W (1998) Control of cyclin ubiquitination by CDK-regulated binding of Hct1 to the anaphase promoting complex. Science 282: 1721–1724

Zhou Y, Ching YP, Chun AC, Jin DY (2003) Nuclear localization of the cell cycle regulator CDH1 and its regulation by phosphorylation. Journal of Biological Chemistry 278: 12530–12536

